# A Game of Thrones at Human Centromeres I. Multifarious structure necessitates a new molecular/evolutionary model

**DOI:** 10.1101/731430

**Authors:** William R. Rice

## Abstract

Human centromeres form over arrays of tandemly repeated DNA that are exceptionally complex (repeats of repeats) and long (spanning up to 8 Mbp). They also have an exceptionally rapid rate of evolution. The generally accepted model for the expansion/contraction, homogenization and evolution of human centromeric repeat arrays is a generic model for the evolution of satellite DNA that is based on unequal crossing over between sister chromatids. This selectively neutral model predicts that the sequences of centromeric repeat units will be effectively random and lack functional constraint. Here I used shotgun PacBio SMRT reads from a homozygous human fetal genome (female) to determine and compare the consensus sequences (and levels of intra-array variation) for the active centromeric repeats of all the chromosomes. To include the Y chromosome using the same technology, I used the same type of reads from a diploid male. I found many different forms and levels of conserved structure that are not predicted by –and sometimes contradictory to– the unequal crossing over model. Much of this structure is based on spatial organization of three types of ~170 bp monomeric repeat units that are predicted to influence centromere strength (i.e., the level of outer kinetochore proteins): one with a protein-binding sequence at its 5’ end (a 17 bp b-box that binds CENP-B), a second that is identical to the first except that the b-box is mutated so that it no longer binds CENP-B, and a third lacking a b-box but containing a 19 bp conserved “n-box” sequence near its 5’ end. The frequency and organization of these monomer types change markedly as the number of monomers per repeat unit increases, and also differs between inactive and active arrays. Active arrays are also much longer than flanking, inactive arrays, and far longer than required for cellular functioning. The diverse forms of structure motivate a new hypothesis for the lifecycle of human centromeric sequences. These multifarious levels of structures, and other lines of evidence, collectively indicate that a new model is needed to explain the form, function, expansion/contraction, homogenization and rapid evolution of centromeric sequences.

## Introduction

The genomic regions that form human centromeres (illustrated in Figure 1) are unusual for several reasons. They are exceptionally large (typical array lengths of 2-3 Mb, but can be as large as 8 Mb; Willard 1991; Miga et al. 2014) and composed of long, complex repeats (repeats of repeats called **H**igher **O**rder **R**epeats [HORs], reviewed in Schueler and Sullivan 2006; McNulty and Sullivan 2018; Willard and Waye 1987) with high levels of homogeneity. They are also defined epigenetically by a high concentration of **CEN**tromere **P**rotein-**A** (CENP-A, a histone H3 variant, aka CEN-H3) rather than by their DNA sequence (reviewed in Black and Bassett 2008). This sequence-independence has led to the sporadic occurrence of ectopic functional centromeres at diverse sites located throughout the single-copy euchromatin (Marshall et al. 2008). Centromeric sequences and their flanking regions also show extreme linkage disequilibrium for SNPs, indicating an exceptionally low rate of crossing over (mitotic and meiotic) between homologs (Roizes 2006; Pironon et al. 2010; Talbert and Henikoff 2010; Langley et al. 2018). The most unique evolutionary feature of centromeres is their exceptionally rapid rate of sequence turnover. This speed is illustrated by the high sequence divergence found between the centromeric repeats of humans and chimps: so high that most fluorescent DNA probes designed to hybridize to human centromeric repeats fail to recognize (even weakly, and at low stringency) the centromeric repeats on their chimp orthologs (Archidiacono et al. 1995). This rapid divergence at centromeric repeats occurs despite the fact that the base substitution rate between these species averages only 1.2% in repeated and non-repeated DNA across non-centromeric regions (Brittan 2002).

**Figure 1.**
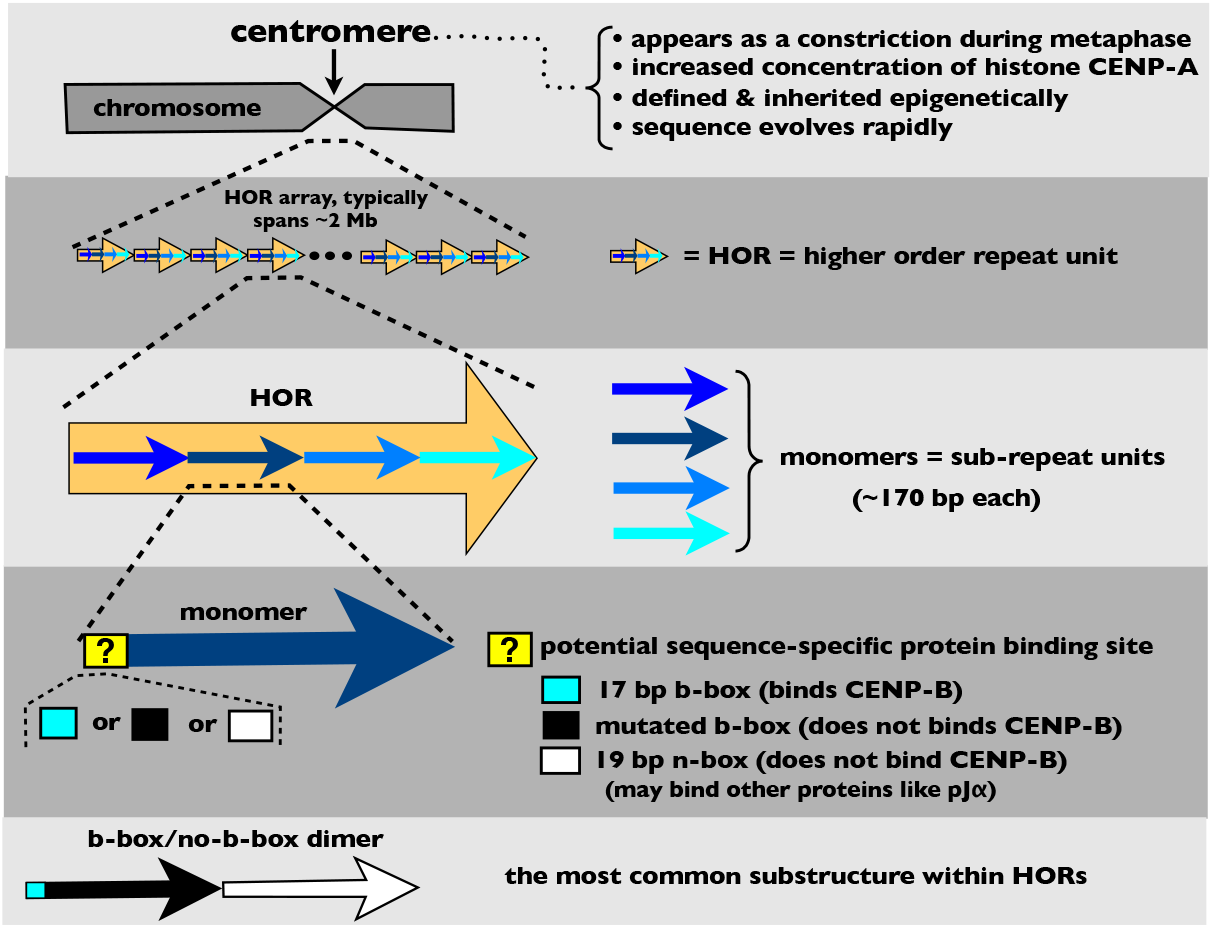
Summary of the structure of human centromeres and **H**igher **O**rder **R**epeat (HOR) arrays.

What molecular mechanism(s) underlies the complex repeat organization, homogeneity, extreme array size, and rapid evolution of human centromeric sequences? The generally accepted model for the organization and evolution of human centromeres was developed by Smith (1976) to explain the evolution of eukaryotic satellite DNA. He showed by computer simulation that mitotic, out-of-register crossover between sister chromatids can generate complex, highly homogeneous repeats that span long stretches of DNA. Stephan (1989) and Stephan and Cho (1994) expanded on this work by showing that the critical parameters are: 1) the ratio of the mitotic recombination rate between sister chromatids (*r*) to the base pair mutation rate (*u*), and 2) the minimum match length (*m*) required to permit out-of-register recombination. For a fixed value of *m*, low values of *r/u* lead to no repetitive structure, intermediate levels generate long repeats, and higher values lead to short repeats. By varying the values of *u*, *r*, and *m*, repeats of a wide diversity of length and complexity can be generated.

Uncertainty in the values of *r*, *u*, and *m* at centromeric DNA, coupled with the wide diversity of repeat structures that can be generated by the Smith model, make this model compatible with the complex repeats observed at the centromeres of human chromosomes –thus making it difficult with current information to disprove the Smith model in the context of centromeric repeats. However, the Smith model predicts that the sequences of centromeres are essentially random (excepting intrinsically harmful sequences, such as those producing undesirable secondary structures, e.g, fold back loops that stall replication forks) rather than being highly structured, as would occur if there were strong functional constraints on repeat sequence. The Smith model also ignores the length-eroding influence of the **S**ingle **S**trand **A**neling (SSA) pathway during the repair of **D**ouble **S**trand **B**reaks (DSBs; Paques and Haber 1999), which had not been discovered at the time of Smith’s paper.

Here I compared all of the the active centromeric sequences across a single human genome (i.e., across one haplotype) to search for intrinsic structure, at multiple levels, that is not predicted by the Smith model. I found many levels of structure at centromeric repeats. This diverse structure demonstrates the need for a new hypothesis for the process(es) that: i) homogenizes repeat sequences within a centromeric repeat array, ii) causes HOR organization to change with increasing length (number of monomers per repeat unit), and iii) generates length variation among homologous repeat arrays (as an alternative to mitotic unequal crossing over between sister chromatids that is assumed by the Smith model). Lastly, the structure that I observed at centromeric repeats motivates a new hypothesis for their lifecycle.

## Searching for all active centromeric repeat arrays within a single genome

The fundamental unit of all established repeats that make up the centromeric sequences of human chromosomes is a monomer (Figure 1) that is ~170 bp in length (Choo et al. 1991). At active centromeres, these monomers can vary in sequence by as much as 35% within and among chromosomes (Fukagawa and Earnshaw 2014). Tandem arrays of centromeric repeats that have been characterized to date contain hundreds to thousands of repetitions of complex repeated units (Figure 1), each of which is made up of between two and 34 different monomers: so they are repeats of repeats and hence classified as **H**igher **O**rder **R**epeats (HORs; Willard and Waye 1987). Based largely on decades of work by Huntington Willard and collaborators, one or more HORs have been mapped to each of the 23 human chromosomes. Reference models for these sequences (stochastically generated sequences of HOR arrays that reflect the sequence diversity and imperfect repeat ordering found among shotgun Sanger reads from the human genome project) can be found at the UCSC genome browser (GRCh38; Miga et al. 2014). Some HORs are shared among non-homologous autosomes, e.g., chromosome groups (1, 5, and 19), (13 and 21) and (14 and 22) and each share a unique HOR in common. At least half of the chromosomes have more than one HOR present simultaneously (UCSC genome browser, GRCh38; Ziccardi et al. 2016). On some chromosomes, more than one HOR is active (Maloney et al. 2012; McNulty and Sullivan 2018), although current evidence indicates that only one HOR is active at one time on a single chromosome (e.g., Aldrup-MacDonaldet al. 2016).

Henikoff et al. (2015) recently identified and quantified the most common HOR monomers within a single human genome and argued that because the functional HORs at each chromosome are embedded within megabases of flanking repeats, there is considerable uncertainty about the sequence of the functioning centromeric DNA on most human chromosomes. Another factor contributing to uncertainty of centromeric sequences is the small number of PCR-generated sequences used to characterize the HORs on most chromosomes. To circumvent this uncertainty, I set out to find –and verify as active– the large-sample-size consensus sequences of all functioning centromeric HORs across all human chromosomes. I did this by analyzing long PacBio SMRT (**Pac**ific **Bio**sciences **S**ingle **M**olecule **R**eal **T**ime) reads from a single homozygous genome. Homozygosity removes ambiguity produced by differing centromeric sequences between homologs. Long reads (up to 40 kb) allowed me to look for structure within long stretches of tandem HORs and ascertain the degree to which monomer number and ordering varied within localized centromeric regions. The long reads also generated hundreds of copies of each monomer that were used for the calculation of their consensus sequence. Finally, I searched this full set of active centromeric HORs for conserved structure (that would not be predicted by the Smith model) at multiple levels.

### Protocol used to find the HOR consensus sequences for all chromosomes within a single haploid genome

The overall strategy was to find long DNA sequencing reads (archived from published studies) that contained short, diagnostic sequences (b-boxes, see below) indicating that they contained HORs that feasibly coded for centromeric sequences. These reads were next cut into small pieces (by cutting immediately before b-box sequences) that each contained a subunit of the HOR. The pieces were then clustered into groups that coded for the same subunit and the consensus of each subunit was determined. These subunit sequences were then mapped back onto the original read to determine their ordering within the HOR. The consensus HOR from the single read was then used to find hundreds of additional reads containing the HOR to determine the centromere-wide consensus sequence. ChIP/Seq sequencing reads (archived from published studies) were then used to determine if the HOR was part of an active centromeric sequence. This procedure was repeated until an active HOR was found for each chromosome within a single haploid genome of a human. These steps are shown in a summary figure found at the end of this section (Supplemental Figure S9).

I started with an SRA (**S**equence **R**ead **Ar**chive) collection of DNA sequence read files available on the NCBI (**N**ational **C**enter for **B**iotechnology **I** nformation) web site (https://trace.ncbi.nlm.nih.gov/Traces/sra/sra.cgi?). The files (SRX533609, located at https://www.ncbi.nlm.nih.gov/sra/?term=SRX533609) were generated using third generation, PacBio SMRT sequence technology that generates long reads but with a high error rate (11-15%, mostly indels; Rhoads and Au 2015). The reads were generated by Chaisson et al. (2015) from a homozygous human female hydatidiform mole (average read length = 5.8 kb and maximum lengths of ~40 kb). I began with a composite file composed of a haphazardly selected collection of SRA files containing 3.25 x 10^6^ reads. This file size will rarely contain multiple copies of single or low copy-number DNA sequences, but should be enriched with many copies of those from long, repetitive sequences. I next used BLAST (**B**asic **L**ocal **Al**ignment **S**earch **Tool**) to search for sequences containing multiple hits closely matching the 17 bp CENP-B-binding consensus sequence (hereafter a ‘b-box’; 5’-Y**TTCG**TTGG**A**AU**CGGG**A-3’, Supplemental Figure S1). I chose this DNA sequence because it binds the only known centromeric protein that has sequence specific DNA binding (CENP-B, which binds when the boldface bases are present and properly spaced; Kipling and Warburton 1997), and because it was previously found to be widespread in HORs from many chromosomes (Masumoto et al. 1989) – but realizing that not all previously reported centromeric HORs contained the sequence (e.g., the only HOR on the Y chromosome). Although this procedure is not guaranteed to find all of the active centromeric HORs, it was expected to find many of them (Masumoto et al. 1989), and possibly all of them except the HOR on the Y chromosome (which is absent in the female hydatidiform mole).

I next haphazardly selected a single, long (> 15 kb), b-box-dense read and divided it into pieces by cutting immediately before each b-box. Pieces were next clustered by constructing a neighborjoining tree and the consensus sequence of each cluster was determined using CLC Sequence Viewer version 7 (= low-N consensus). A typical cluster of a PacBio read cut at b-boxes is shown in Supplemental Figure S2). I next BLASTed the low-N consensus sequences of the pieces back onto the original read to determine their consensus order. I then BLASTed these low-N consensus sequences (in pieces of 2-3 contiguous monomers) back on to the PacBio hydatidiform mole reads (SRX533609) to obtain > 300 hits and took the consensus of these DNA sequences to form a large-N consensus of each piece. Finally, I BLASTed the large-N consensus pieces onto the human genome sequence (GRCh38) to determine which, if any, chromosome(s) were known to contain the sequences at high copy number (on the reference model sequences of their centromeric HORs). I repeated these steps until I found at least one consensus HOR for each autosome and the X.

To find the centromeric sequences for the Y chromosome based on long PacBio SMRT reads, I used a publicly available SRA file from a diploid male (AK1; https://www.ncbi.nlm.nih.gov//bioproject/PRJNA298944; Seo et al. 2016). I used BLAST to search for PacBio SMRT reads containing any of the 34 monomers previously found on the human Y chromosome and included within the reference model for this chromosome in GRCh38 (Miga et al. 2014). Using these reads as a data base, I calculated a consensus for each of the 34 monomers (using at least 40 monomer copies) and confirmed that the order of monomers was the same as that shown in the Y chromosome reference model in GRCh38 (Miga et al. 2014).

### The consensus HOR sequences for all chromosomes

The steps described in the above section generated 19 large-N consensus HORs that collectively mapped to all of the autosomes or the X (some mapped to more than one chromosome) plus an additional sequence for the Y chromosome from the diploid male genome (total HORs = 19 + 1 = 20). The consensus sequences for the 20 identified HORs are listed in Supplementary Table S1. Note that one HOR maps to chromosomes 1, 5, and 19, a second HOR to chromosomes 13 and 21, and a third HOR to chromosomes 14 and 22 (see Ziccardi et al. 2016 for the logic used to resolve HORs located on the 13/21 and 14/22 groups). The fact that three groups of chromosomes each share a different HOR indicates that each group is feasibly undergoing concerted evolution. Some process, such as ectopic gene conversion, would have to generate sufficient exchange among non-homologous chromosomes to maintain this sequence homogenization.

In Supplementary Table S1 (and all later figures based on this table) I show the predominant consensus HOR sequence for each chromosome. For most chromosomes, the large majority of long reads that I examined were highly homogeneous and matched the consensus HOR sequence closely. The HORs on some chromosomes, however, were heterogeneous. The PacBio reads with HORs mapping to chromosome 1 (which also mapped to chromosomes 5 and 19) were exceptionally variable. The predominant repeat was the simple dimer (a 2-monomer HOR; one monomer with, and one monomer without a b-box; see bottom of Figure 1) listed in Supplementary Table S1, but other b-box/no-b-box dimers were also present. I found 4 clusters of b-box/no-b-box dimeric sequences (Supplemental Figure S3). In a sample of 510 PacBio reads with a total of 19.5 x 10^3^ dimers that mapped to chromosome 1, 66.1% of the dimers matched closely to the sequence shown in Supplemental Table 1 (which I will refer to as dimer-1), and about a quarter of the reads were a contiguous repetition of this dimer over >10 kb (Supplemental Figure S4-A). However, most reads were a heterogeneous mixture of mostly dimer-1 but also containing one or more of three other dimers (e.g., see supplemental Figure S4-B). All of the dimers found on chromosome 1 shared at least 90% sequence similarity with dimers found on the longer (6-dimer) HOR located on chromosome 16 –a pattern I come back to in a later section. Interestingly, ~3% of the reads were homogeneous (or nearly so) for a 4-dimer HOR (Supplemental Figure S4-C).

On chromosome 2, most of the HORs were composed of the 2-dimer HOR shown in Supplemental Table 1 (each dimer contains a b-box monomer and a no-b-box monomer), but while some long reads were perfectly homogeneous (Supplemental Figure S5-A), most had numerous imperfections in which the same dimer was tandemly repeated two or a few times (Supplemental Figure S5-B). On chromosome 9, the consensus HOR shown in Supplemental Table 1 (a 15 monomer HOR composed of 7 b-box/no-b-box dimers and one lone no-b-box monomer) was the predominant repeat, but while some long reads were homogeneous for this HOR (Supplemental Figure S6-A), most long reads contained repeats with indels (one or more missing/added dimers) interspersed with the consensus repeat (Supplemental Figure S6-B). Chromosomes 4, 8, and 15 had a pattern similar to that seen on chromosome 9. In the following analyses, I will focus on the consensus sequences for each chromosome that is shown in Supplemental Table 1.

### Confirming that the consensus HOR sequences were active centromeres

To confirm that the sequences that I found were the active centromeric HORs, I used SRA files from a ChIP-seq study of human centromeres from the HuRef lymphoblastoid line (Illumina 100 bp paired-end reads from the genome of J. C. Venter; SRA file GSE60951; https://www.ncbi.nlm.nih.gov/bioproject/?term=GSE60951; CHiP based on CENP-A; Henikoff et al. 2015). I BLASTed a large number of Illumina 100 x 100 bp paired-end reads (4.76 x 10^6^, from either the ChIP or Input SRA files) onto each of the 20 large-N consensus HOR sequences. I recorded the location of each BLAST hit (to reduce off-target hits, they needed to be at least 97 bp long and match at least 97% of base positions) and calculated the ChIP and reference hit rates for each monomer of each of the 20 large-N HOR consensus sequences. As a control, I used the consensus sequence of a flanking HOR on chromosome 15 (GJ212851.1 in UCSC genome browser [GRCh38]) which has not been found to be active in past CHIP/Seq studies (e.g., Nechemia-Arbely et al. 2017; see also Supplemental Figure S7 adapted from Nechemia-Arbely et al. 2017). All large-N consensus HOR sequences were found to have a high ChIP-seq ratio, compared to the inactive control, indicating all were the active repeats of centromeres (Supplemental Figure S8). As a second check on the activity of the HORs that I identified from PacBio reads, I used CHiP-seq data (CHiP based on CENP-A) from Nechemia-Arbely et al. (2017) on a different cell line (HeLa). The centromere reference models corresponding to each of the HORs that I report in Supplementary Table S1 show strong CENP-A enrichment –indicating an active status for each of these sequences (Supplemental Figure S7). A summary of the step-by-step procedures that I used to to find –and verify as active– the 20 consensus HORs are tabulated in Supplemental Figure S9.

## Multifarious structure at centromeric HOR arrays

### Head-to tail orientation of monomers is completely conserved

All of the monomers from all of the active, centromeric HORs can be aligned to the consensus HOR sequence across human chromosomes reported by Choo et al. (1991). This alignment allows the 5’ and 3’ end of each monomer to be identified. In past studies, a head-to-tail orientation (i.e., 5’ to 3’) of all monomers within HORs has been reported (e.g., Waye et al. 1987a,b; Haaf and Willard 1992). In the genomewide screen reported here (Supplementary Table S1, and illustrated graphically in a latter section by Figure 4), every monomer of every consensus, centromeric HOR has consistent head-to-tail orientation. This strong structural pattern across all 20 centromeric HORs would be unexpected if the HORs were evolving via unequal crossing over between sister chromatids because inversions among the constituent monomers of HORs would be neutral and therefore expect to accumulate by drift within different lineages of the centromeric HORs.

### Spacing of b-boxes is highly non-random, generating a predominance of b-box/no-b-box dimers

To look for additional structure among the full genomic complement of consensus, centromeric HORs, I examined the distribution of distances separating b-boxes (including ‘broken b-boxes’ with mutations predicted to block the binding of the CENP-B protein; Figure 2). The units in Figure 2 are the number of monomers lacking b-boxes separating monomers that contain b-boxes. I excluded from this analysis HORs from the male-limited Y chromosome which is devoid of b-boxes. Multiple b-boxes were never found within a single monomer and the greatest number of no-b-box monomers separating sequential b-box monomers was two. By far the most common (81%) spacing was one b-box every other monomer (i.e., one no-b-box monomer separating b-box monomers). If b-box and no-b-box monomers were arranged randomly, the spacing between b-boxes (in units of number of no-b-box monomers separating b-box monomers) would follow a negative binomial distribution and the observed spacing strongly deviates from this null expectation (Figure 2; p = 5.27 x 10^-51^, Chi-square goodness of fit test). There are far too many b-box monomers separated by one no-b-box monomer and far too few separated by zero or > 1 monomers. This far-from-random pattern demonstrates that the predominant substructure of the active centromeric HORs is a dimer made up of b-box- and no-b-box-monomers. I will refer to this predominant HOR substructure as a b-box/ no-b-box dimer (see bottom of Figure 1).

**Figure 2.**
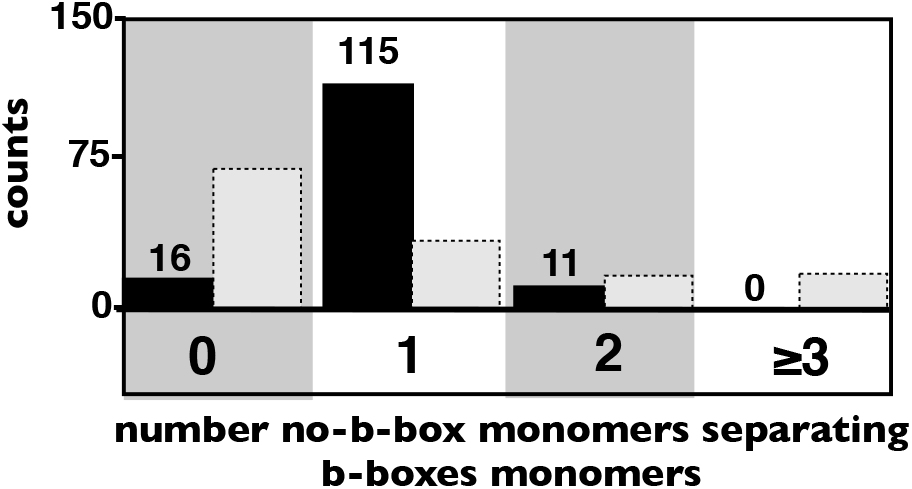
The distribution of distances separating CENP-B-boxes within the 19 active consensus HOR sequences from the hydatidiform mole (CHM1). Observed values are dark bars and random expectations (from negative binomial distribution) are light bars. The X-axis units are monomers (~170 bp).

Based on a much smaller sample of HORs, Thomas et al. (1989) also reported a strong tendency for monomers to be organized as dimers within human centromeric repeats. Monomers in this study were classified based on monomer subfamily types. Some authors appear to claim that all human alpha satellite DNA is arranged as b-box/no-b-box dimers (e.g., Tawaramoto et al. 2003), but the HOR sequences reported here for all chromosomes, and those reported previously for individual chromosomes (e.g., Durfy and Willard 1989; Waye and Willard 1986b), do not support this claim.

### Both monomers within b-box/no-b-box dimers have conserved sequences at their 5’ ends

I next aligned (CLC Sequence Viewer version 7) all b-box monomers, and separately all no-b-box monomers, from all b-box/no-b-box dimers that contained a functional b-box (i.e., one predicted to bind CENP-B; Masumoto et al. 1989; Kipling and Warburton 1997). Although my search for functional b-box monomers only required their b-box to match the 9 base pairs needed for the binding of the CENP-B protein (minimal b-box = -TTCG----A--CGGG-; Masumoto et al. 1989; Kipling and Warburton 1997), I found strong conservation for the all but the 1st position, which as predicted by the consensus sequence, was nearly always C or T (see heat map in Supplemental Figure S10A). This strong conservation at nearly all of the 17 bp positions of a b-box indicates positions outside the minimal b-box –although not absolutely required for CENP-B binding– may nonetheless facilitate the binding of this protein (or have some other selectively-favored function).

A heat map of sequence conservation for no-b-box monomers (from all b-box/no-b-box dimers that bind CENP-B) showed a different highly conserved region (spanning 19 bp) near the 5’ end of these monomers. It is displaced 6 bp downstream from the corresponding 5’ start site of the b-box in b-box monomers (Supplemental Figure S10B). The consensus sequence for this “n-box” (an abridgment of no-b-box) is T[A/G][A/ G]AAAAGGAAATATCTTC. Using a smaller sample of centromeric repeats, Gaff et al. (1994) and Romanova et al. (1996) also reported a conserved region on no-b-box monomers that binds the pJα protein (a pJα-box, aka an a-box), but this 17 bp sequence started at the corresponding 5’ position (on no-b-box monomers) as the b-box on b-box monomers. In my larger sample of centromeric repeats with confirmed centromeric functioning, I found only weak sequence conservation for the first 6 bp of the pJα-box, but I found strong sequence conservation extending 8 bp beyond its 3’ boundary (i.e., spanning the 19 bp n-box region; Supplementary Figure S10A,B). The n-box reported here is also substantially conserved in the 34 no-b-box monomers that make up the consensus sequence of the active HOR on the Y chromosome (Supplemental Figure S10C). The consensus sequence for the 5’ end of n-box monomers also contains all of the properly positioned bases required for the binding of the pJα protein (Gaff et al. 1994), but this study did not identify the minimal sequences needed for the binding of the pJα protein. At this point forward, I will define the subset of b-box/no-b-box dimers that carry a functional b-box (i.e., one predicted to bind CENP-B) to be *canonical b-box/n-box dimers*, which I will abridge to b/n-box dimers. When the b-box is mutated so that it is predicted to no longer bind CENP-B (i.e., it does not contain the minimal b-box sequence), I refer to them as *broken b/n-box dimers*. I will also use the term *broken b-box* to be a sequence nearly identical to the b-box consensus, but not predicted to bind CENP-B.

### b-boxes at every other nucleosome may be common in other mammals in which CENP-B binds centromeric repeats

Previous studies in humans indicate that the b/n-box dimeric structure strongly influences the positioning of nucleosomes (Tanaka et al. 2005; Hasson et al. 2013; Fujita et al. 2015; Henikoff et al. 2015). Each dimer winds around two neighboring histone cores leading to an alternating pattern of linkers between nucleosomes: one with a b-box and the next with an n-box (see Supplemental Figure S12 in the companion paper [Rice 2019] where I will discuss the potential significance of this alternation pattern).

The finding that most monomers at human chromosomes are organized as b/n-box dimers (which are about 170 + 170 = 340 bp long, Figure 2), motivated a search to see if this configuration is common in other mammals that have evolved b-boxes at their centromeric repeats. In chimps, sequences of centromeric HORs on orthologous chromosomes are only distantly related to those of humans (Archidiacono et al. 1995). Despite this large-scale sequence divergence, the distribution of spacing between b-boxes in long PacBio reads is nearly identical between humans and chimps, with a strong mode near 340 bp (Supplemental Figure S11). This similarity suggests that the predominant b-box/no-b-box dimeric arrangement seen in humans has persisted in both species despite substantial sequence divergence of the orthologous centromeric HORs.

In the more distantly related house mouse (*Mus musculus*), centromeric monomers on the X and autosomes are found in the minor satellite. These monomers are not organized into well defined HORs and all monomers contain a b-box-like 17 bp sequence –the consensus of which exactly matches human consensus except at the first position, where A replaces T (Wong and Rattner 1988; Masumoto et al. 1989; Kipling et al. 1994; Komissarov et al. 2011). However, work by Iwata-Otsubo et al. (2017) found that a substantial proportion of these b-boxes did not match the minimal sequence needed to bind CENP-B (Kipling and Warburton 1997). In a large random sample of mouse minor satellite monomers, I found that about half have b-box sequences that are not predicted to bind CENP-B, and that these appear to be randomly distributed throughout the centromeric repeat arrays (Supplemental Figure 12). The high frequency of nonfunctional b-boxes, their random distribution, and their positioning within nucleosomes (Iwata-Otsubo et al. 2017) would be expected to plausibly make it common for neighboring linkers between nucleosomes to alternate (one with and one without CENP-B binding): matching the predominant pattern in humans.

In a diverse sample of new world monkeys, centromeric monomers were found to be ~345 bp long (range = 340-350), and b-boxes have apparently evolved independently in three species (marmosets, squirrel monkeys and tamarins; Kugou et al. 2016). In each species only a single b-box per monomer has evolved, leading to ~340 bp b-box spacing that is very similar to the predominant spacing found in humans. Centromeric b-boxes that bind CENP-B have also been reported in red-necked Wallabies (Bulazel et al. 2006), green monkeys (Yoda et al. 1996) and also tree shrews, giant pandas, gerbils, and ferrets (see Kipling and Warburton 1997): but too little sequence is available to determine the distribution of spacing between functional b-boxes in these species.

Collectively, these studies in mammals indicate that the positioning of b-boxes at every other linker between nucleosomes (the predominant pattern in humans) may be common. This pattern across distantly related species suggests that the juxtaposition of nucleosomes with and without CENP-B-binding may have functional significance. A hypothesis explored in the companion paper (Rice 2019) is that CENP-B bound to linkers between nucleosomes strengthens kinetochores by recruiting more CENP-C (Fachinetti et al. 2015), but when the binding of this large protein (a dimer with 80-kDa subunits) is not frequently interrupted by CENP-B-free linkers, it interferes with other necessary centromeric molecular interactions, such as the transcription by RNA-Pol-II (Scott 2013; McNulty et al. 2017).

### Centromeric n-box and b-box monomers differ in consensus sequence, length, and variation characteristics

The consensus sequences of n- and b-box monomers from canonical b/n-box dimers are diverged by 15.8% and most of this divergence (70%) is outside the n- and b-box regions (Supplemental Figure S13). The n- and b-box monomers also differ in average length (170.8 vs 168.7 bp, respectively, p < 0.0001, Welsh test) and b-box monomers are far more variable in length (Var[b-box-monomer] / Var [n-box-monomer] = 8.6, p < 0.0001, Levine test). The differences in length parameters suggest that there may be different functional constraints on the size of the two major monomer types.

### A graphical classification of monomers and dimers within human HORs

To further search the 19 HORs from the hydatidiform mole for structural characteristics, I expressed each HOR symbolically. The idea here is that replacing DNA sequences by symbols of their internal structure would generate insight into their organization: much like replacing an ordered list of the serial numbers of train cars (an array of non-informative numbers) by symbols of their internal structure (engine, caboose, box car, tanker car, etc.) would provide more insight in the structure of trains.

To generate a schematic representation for the the the sequences of the monomers and dimers constituting each HOR (Supplementary Table S1), I began by a cluster analysis (neighbor-joining tree) of all the monomers. They clustered into three well-defined groups: b-monomers, n-monomers, and intermediate monomers. Nearly all b-monomers had a b-box at their 5‘ end and while nearly all n-monomers had an n-box at this location. However, the n- and b-box sequences alone did not determine cluster identity. For example, some monomers with a b-box clustered with the n-monomers (examples are shown in Supplemental Figure S14). Dimers were also classified by how much they deviated from a canonical b/n-box dimer, as described in the next paragraph.

In Figure 3, monomers are depicted by small arrows: b-monomers are black, n-monomers are white, and those that clustered intermediately between these types are grey. The b-boxes are classified as: i) functional (containing the 9 base pairs with the proper spacing required to bind the CENP-B protein, depicted by a blue box at the left side of a small arrow), or ii) non-functional (black box at the left side of a small arrow). Each dimer (large arrows bracketing a b-monomer followed by an n-monomer) is classified as: i) canonical (a b/n-box dimer in which there is a CENP-B-binding b-box monomer followed by an n-box monomer; depicted in dark red), ii) broken (mutated b-box that does not bind CENP-B, but otherwise the same as a canonical b/n-dimer; white with dashed outline), iii) non-canonical (b-box spacing is every other monomer but the n- and/or b-box monomer is atypical [see Figure 3]; pink), iv) out-of-context (the monomer preceding a canonical b-box/n-box dimer contains a b-box [data from Hasson et al. 2013 {their Figure 6} indicated these arrangements may interfere with CENP-A recruitment and normal nucleosome positioning]; orange), or v) both non-canonical and out-of-context; yellow). Supplemental Figure S14 illustrates how monomers and dimers were classified in the context of specific HORs. Figure 4 depicts all of the 19 HORs found in the hydatidiform mole (plus the Y HOR from the diploid male genome) using this schematic representation. The figure visually illustrates the substantial predominance b/n-box dimers within the centromeric HORs found on the X and autosomes: a highly non-random structure.

**Figure 3.**
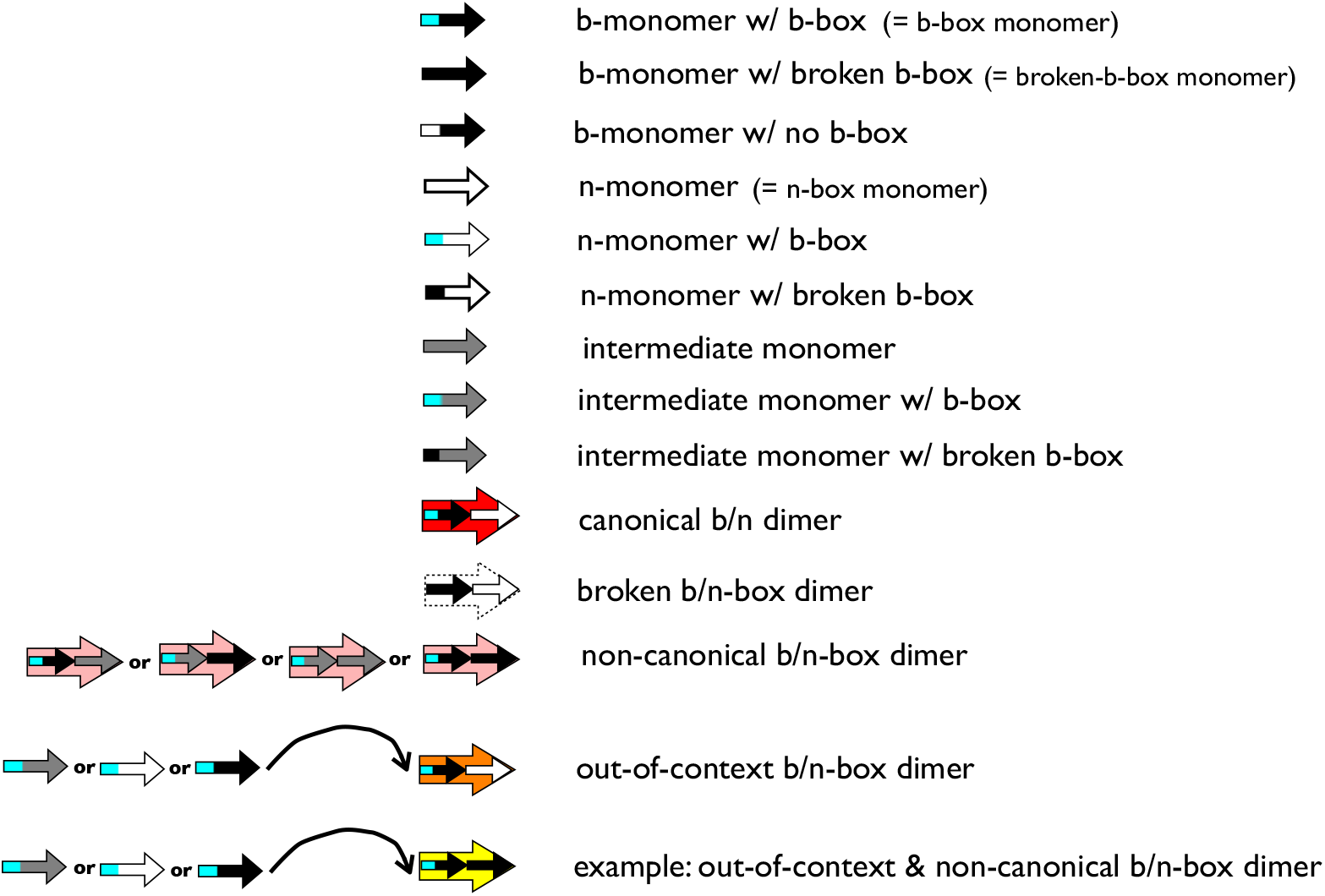
A classification framework for monomers and dimers. Monomers are classified by their cluster position (b-monomer, n-monomer, or intermediate) in a cluster analysis based on their entire sequence.

**Figure 4.**
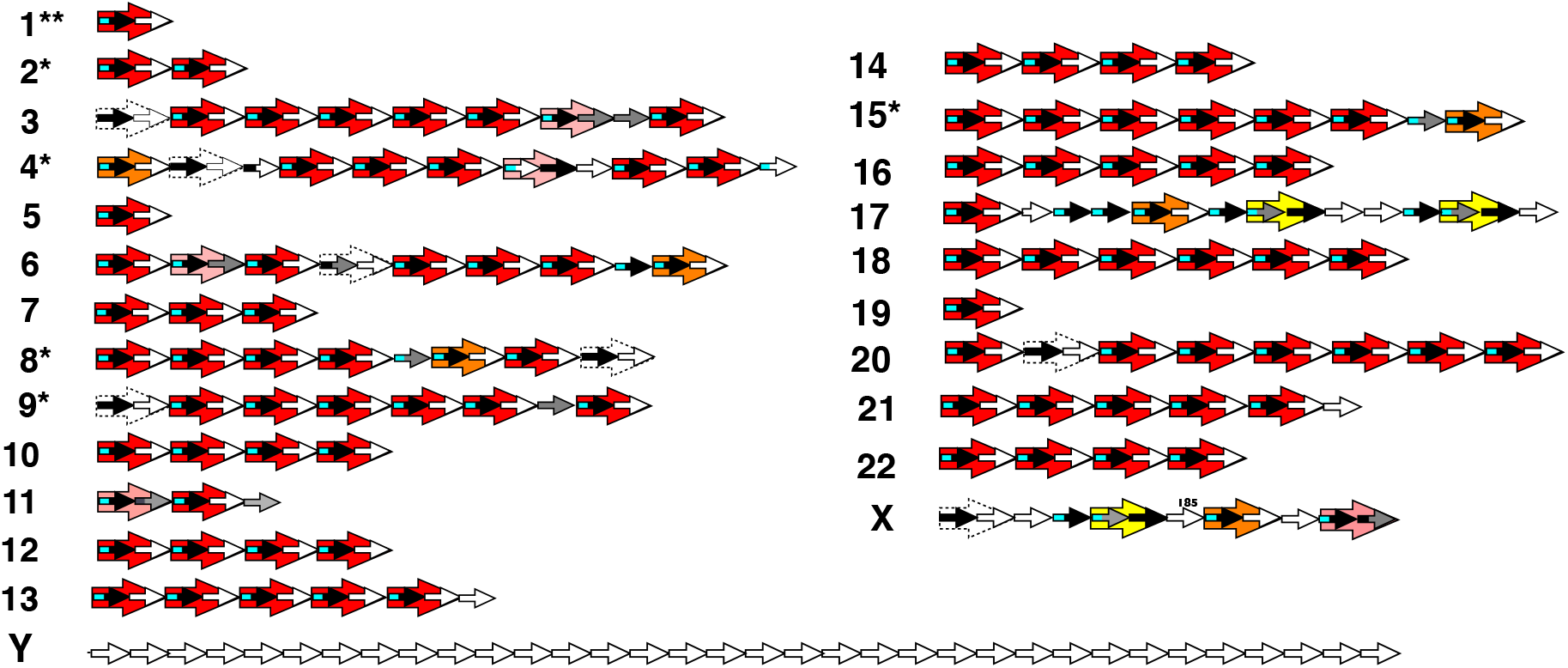
Schematic diagrams of the 19 consensus HORs from from PacBio reads from a hydatidiform mole genome (CHM1), plus the consensus HOR of the Y chromosome from PacBio reads from a diploid male genome (KOREF): see Figure 3 for arrow key. The 185 above an n-box monomer within chromosome X, denotes its unusual length of 185 bp, rather than a value close to 170 bp, as is seen in all other monomers. Long reads from chromosomes with an asterisk have primarily the consensus sequence but indels (of one or more monomers) are not uncommon (see Supplemental Figures 5–6). Chromosome 1 has two asterisks because it is especially variable and contains long reads with up to threa additional mimers (see Supplemental Figure S4). Note that b/n-box dimers within the same HOR (large red arrows) do not have the same sequence but do have the same dimeric structure (b-box monomer + n-box monomer) This is also true for b/n-box dimers in different chromosomes with the exception of three groups of chromosomes ([1, 5, and 19], [13 and 21], and [14 and 22]) where each of the groups shares a group-specific HOR sequence.

### Dimers within an HOR share sequence similarity

To search for additional structure within active centromeric HORs, I next made a neighborjoining tree (CLC Sequence Viewer version 7) for all of the 108 dimers (large arrows) shown in Figure 4. There are four well defined clusters with substantial bootstrap support (Figure 5, Supplemental Figure S15). I named each cluster by the lowest number chromosome in the group (human autosomes are labelled based on size, with 1 = largest and 22 = smallest), with the exception of the cen-6-like cluster which contains all but one dimer from the chromosome 6 cluster. When I clustered all monomers from all HORs, clustering similar to that found among dimers was observed within the n-box and b-box monomer types (Supplemental Figure S16).

**Figure 5.**
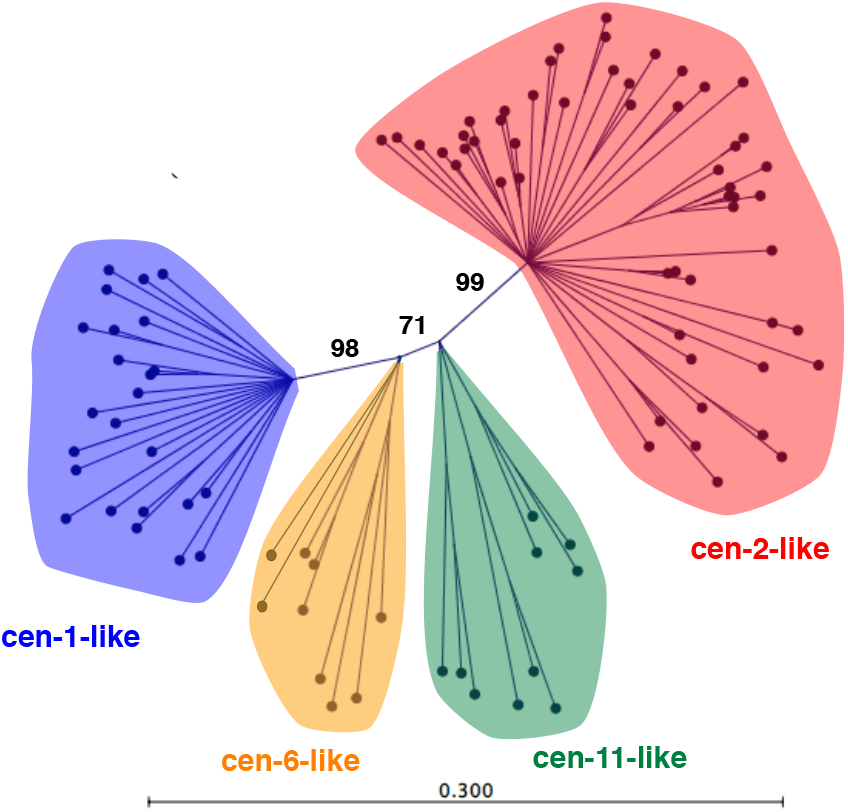
A neighbor-joining tree of the dimers (pairs of bmonomers and n-monomers enclosed by arrows in Figure 4) from all of the consensus HORs from the hydatidiform mole genome (CHM1). Numbers between clusters are bootstrap support values. Lineages with less than 67% bootstrap support are collapsed. Clusters are named by the lowest-number chromosome they contain except the cen-6-like cluster, which is composed entirely of dimers from chromosomes six with a single dimer from chromosome three.

I next looked for structure among the dimers that make up each HOR. In Figure 6, I color-coded all of the dimers from all of the 19 autosomal and X HORs (Figure 4) based on their cluster affiliation in Figure 5. There is a strong pattern: if one dimer in an HOR comes from a cluster then so too do all others, with only two rare exceptions to this rule (< 2%). This pattern suggests that HORs are not random combinations of monomers but groups that share sequence similarity. This pattern indicates that any recruitment of dimers or monomers to an HOR (during their initial establishment or when new repeat elements are added to an established HOR) is facilitated by sequence similarity (Supplemental Figure S17). High sequence similarity, however, is not required. This is illustrated by the HOR shared by chromosomes 14 and 22, where none of the dimers are closely related despite all coming from the same cluster (Supplemental Figure S15).

**Figure 6.**
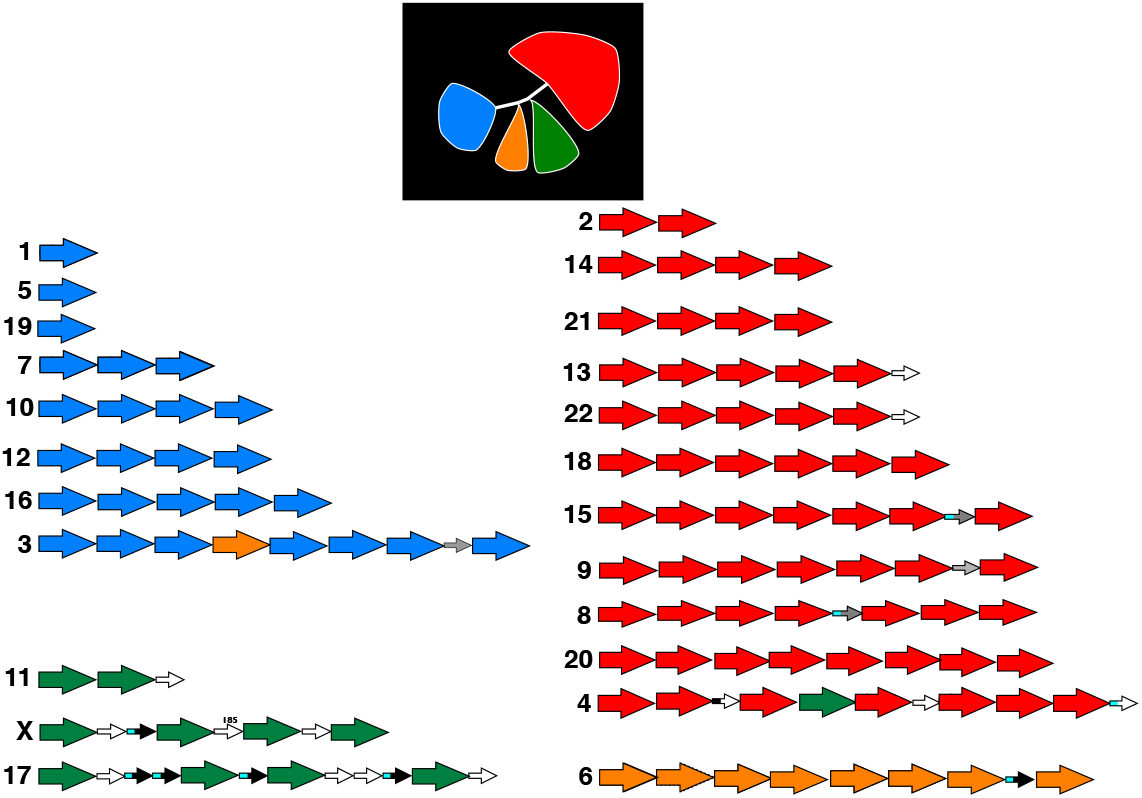
The 19 consensus HORs for the X and autosomes from the hydatidiform mole genome (CHM1) where each b-box/n-box dimer shown in Figure 2 is replaced by a colored arrow with the color matching the dimer cluster to which is was assigned in Figure 5 (summarized in the black square). There is a strong pattern: if one dimer comes from a cluster, then so do all other dimers with only two exceptions to this rule.

### HOR structure changes with HOR length

I also looked for structure among the 20 HORs by arranging them by size (Figure 7). Canonical b/n-box dimers dominate the composition of shorter HORs (≤ 10 monomers in length) whereas larger HORs nearly always have at least some deviations from canonical dimer structure (logistic regression: presence/absence of any monomers outside of canonical dimers vs. number of monomers; p < 0.0001; Kendal’s concordance test: proportion monomers outside of canonical dimers vs. number monomers, p = .0004). I also found that the proportion of b-monomers with broken b-boxes increased with increasing HOR size (Kendal’s concordance test: proportion monomers with broken b-boxes vs. number monomers, p = .0161). In the above tests I excluded the Y chromosome (which has no b-box monomers), but all of the p-values become more statistically significant when the Y chromosome is included. Chromosome 11 is a clear outlier. However, the two b-box/no-b-box dimers and the single lone no-b-box monomer that make up this consensus HOR are closely related to a contiguous piece of one of the flanking, inactive HORs on chromosome 11 (Supplemental Figure S18A): suggesting that this HOR is feasibly a contraction (via a multi-monomer deletion) of a longer flanking (plausibly older) HOR. These observations collectively indicate that HOR structure is dominated by canonical b-box dimers in shorter HORs and that this modular structure is diminished in longer HORs -in substantial part by the reduction of functioning b-boxes.

**Figure 7.**
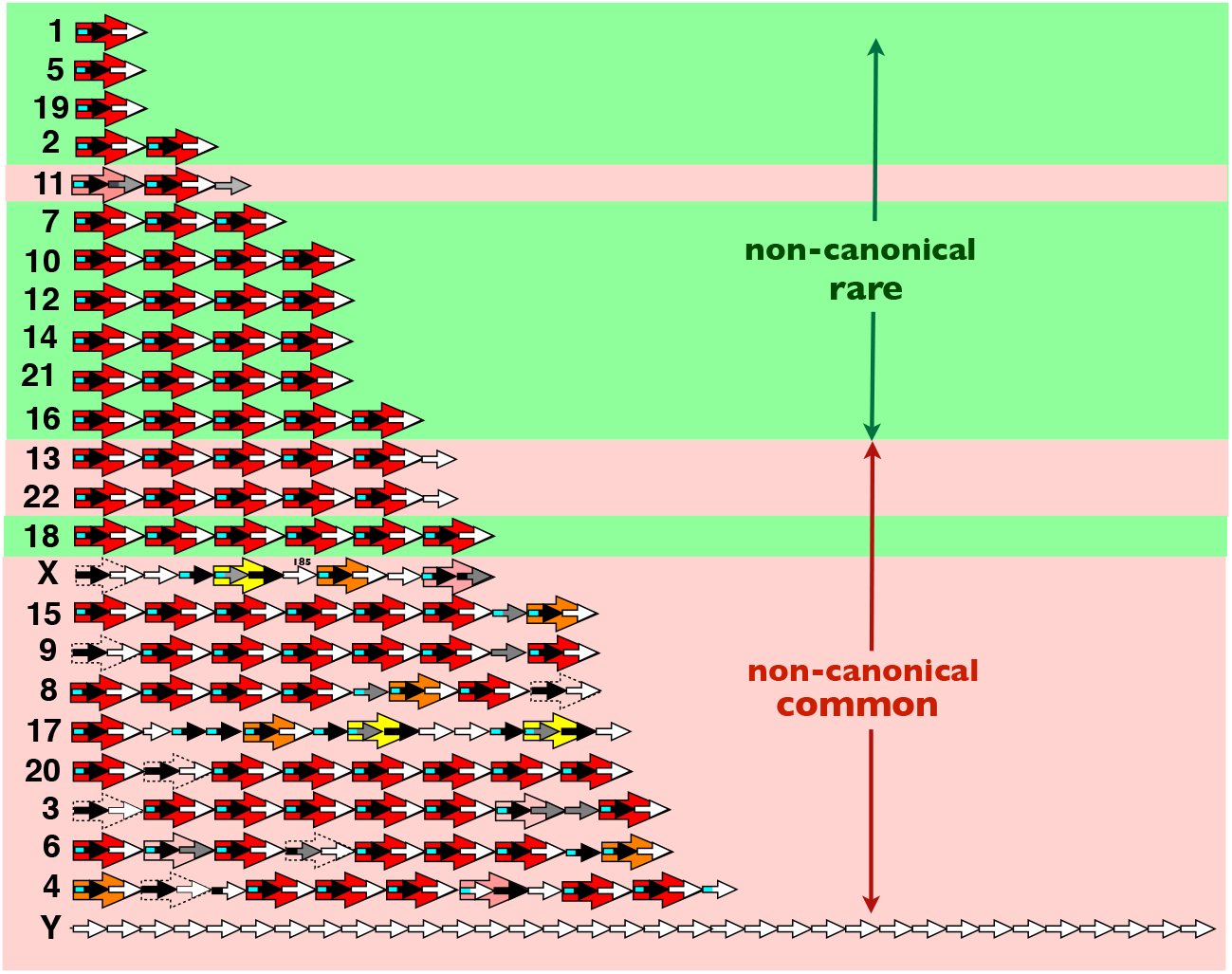
The 19 consensus HORs of the X and autosomes from the hydatidiform mole genome (CHM1) plus the Y sequence from a Korean genome (KOREF) arranged by size. Short HORs are composed almost exclusively of canonical b/n-box dimers, whereas most longer HORs have departures from this modular structure.

Another interesting pattern seen in the ordering of HORs from smallest to largest is that the dimers that make up the shortest HORs are closely related to dimers found in longer HORs on other chromosomes. The one-dimer HOR shared by chromosomes 1, 5, and 19 is 95.7% identical in sequence to the b dimer (second from left in Figures 4 and 7) of chromosome 16 (Supplemental Figure S18B). Similarly, the two dimers of the active HOR on chromosome 2 are closely related to dimers g (second from right; 96.4% identical in sequence) and h (right-most dimer; 93.5% identical in sequence) of the longer active HOR on chromosome 20 (Supplemental Figure S18B). And as mentioned above, the active HOR on chromosome 11 is feasibly a contraction (via a multi-monomer deletion; Supplemental Figure S18A) of a longer flanking HOR (which is inactive). These observations suggest that pieces of HORs can be exchanged among chromosomes and that short HORs are feasibly derived from pieces of longer HORs on different or the same chromosomes.

### Low sequence similarity is common between flanking HORs on the same chromosome

At least 14 of the 24 human chromosomes have one or more HORs that flank the active HOR (UCSC genome browser [GRCh38]). These flanking HORs are usually inactive (e.g., see Supplement Figure S7) but in some cases a flanking HOR may be the active HOR in a minority of individuals (feasibly this occurs when there is a large deletion in the typically active HOR, e.g., on chromosome 17; Aldrup-MacDonald et al. 2016). In Figure 8, I have color-coded the active HORs (circles with dark perimeters and a white central spot) in the HuRef lymphoblastoid line and (from consensus sequences from centromere reference models found in the UCSC genome browser [GRCh38]) the closest flanking HORs (right, left, or both when both are present). Some chromosomes have more than one flanking HOR on one or both sides (UCSC genome browser [GRCh38]) but here I analyze only the closest flanking HORs, because these are predicted to be the youngest (Schueler et al. 2005; Shepelev et al. 2009) and hence least degraded by mutation since they became inactive. Colors depict the cluster (Figure 5) to which the dimers in the HOR map. The yellow cluster represents the 17 “dimers” from the Y chromosome that were made by pairing neighboring monomers (all of which lack b-boxes). The grey cluster contains a group of related dimers only found in the flanking HORs. Flanking arrays on the acrocentric chromosomes 13, 14, 21, and 22 are incompletely resolved on the UCSB genome browser, so I report a single flanking HOR for this group labeled 13^+^. In sharp contrast to the strong cluster-concordance seen for dimers within HORs, flanking HORs are just as likely to be cluster-discordant as cluster-concordant, indicating that sequence similarity has a weaker role in the recruitment of new HORs to chromosomes. If unequal crossing over generated new centromeric HORs, flanking HORs would be derived ‘one from the other,’ and hence they should be closely related: but contrary to this prediction, flanking HORs were commonly distantly related.

**Figure 8.**
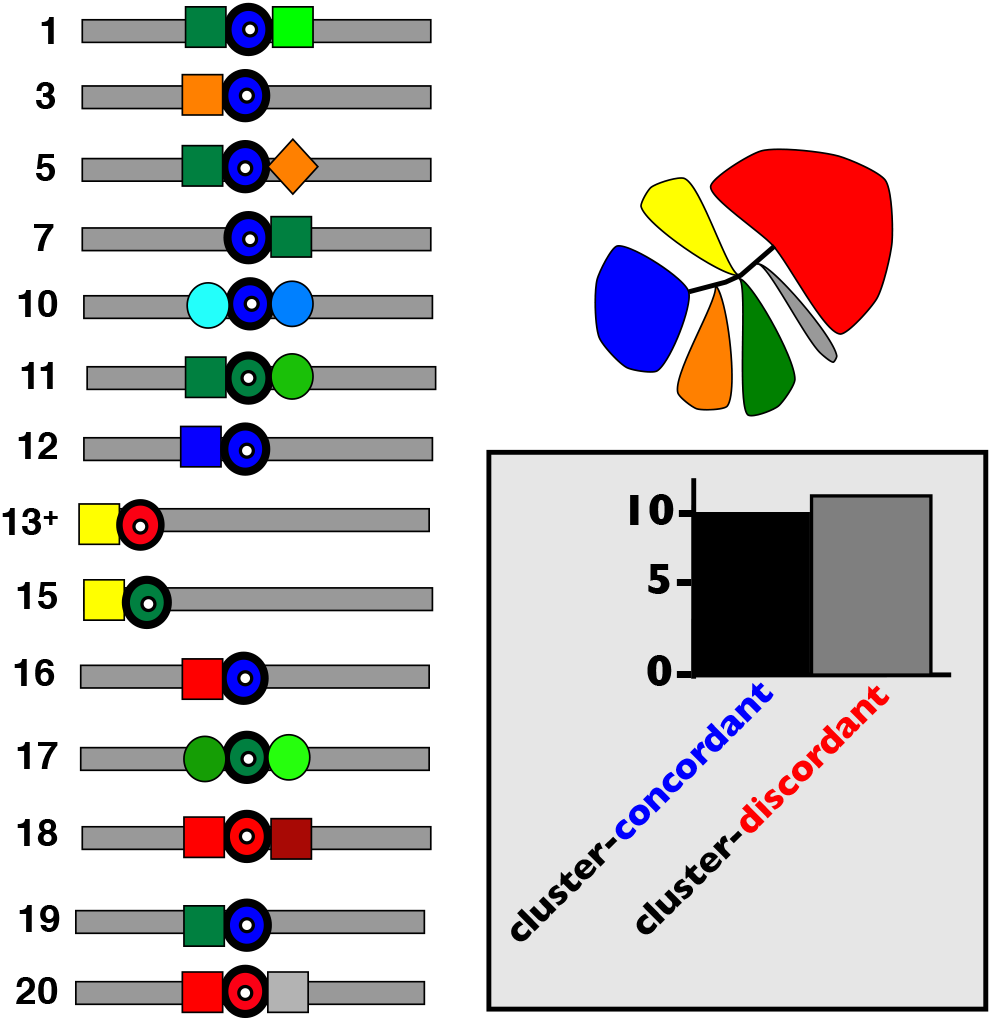
The HORs immediately flanking the active HORs on the chromosomes that contain them (the active HORs are depicted by bold circles with white centers). All HORs are color-coded by the dimer cluster they map to in Figure 5. The grey cluster represents a new cluster (not shown in Figure 5) found only in the flanking HORs and the yellow cluster represents ‘dimers’ on the Y chromosome constructed by pairing adjacent monomers: 1 and 2, 3 and 4, 5 and 6, …, 33 and 34). Flanking HORs on the same chromosome that lack close sequence similarity are depicted by different shapes. Flanking HORs on the same chromosome that show strong sequence similarity are depicted by the same shape but different shades of the same color. Flanking HORs are nearly equally likely to be cluster-concordant with the active HOR as they are to be cluster-discordant.

### Flanking, inactive HORs are smaller and have low canonical b/n-box dimeric structure

Inactive, flanking HOR arrays are generally much smaller than their active counterparts (Supplementary Figure S7) and they have been hypothesized to be both nurseries (i.e., they will expand and become the HORs of the of future) and graveyards (i.e., they are shrinking toward extinction via recurrent deletions). I compared the structure of active HORs (active status indicated by ChIP-seq analysis; Supplemental Figure S7–S8) and inactive flanking HORs (inactive status indicated by ChIP-seq analysis, Supplemental Figure S7). It should be noted that some “inactive”, flanking HORs can (usually at low frequency) be the active centromeric sequence in some individuals (see Table 1 of McNulty and Sullivan 2017). For each of the human chromosomes known to have flanking HORs, I have (in Figure 9) schematically depicted consensus sequences of the active and inactive (immediately flanking) HORs (as described in Figure 4). Consensus sequences for the flanking, inactive HORs were calculated from the reference model sequences found in the UCSC genome browser (GRCh38; Miga et al. 2014). The pattern is visually apparent and born out by statistical analysis: on average flanking HORs are longer (more monomers per repeat unit; Welsh test, p = 0.0123), have a lower density of canonical b-box dimers (Welsh test, p < .0001), have a lower density of monomers that fall into the b-monomers cluster (see Supplemental Figure S14; Welsh test, p < .0001), and have a higher density of monomers with broken b-boxes (that are predicted to not bind CENP-B) (Welsh test, p < 0.0001). In sum, flanking, inactive HORs on average have: i) longer repeats (more monomers), ii) shorter array sizes, iii) reduced canonical b/n-box dimer structure, and iv) more broken b-box monomers (that are predicted not to bind CENP-B). These structural differences are not predicted by the unequal crossing over model.

**Figure 9.**
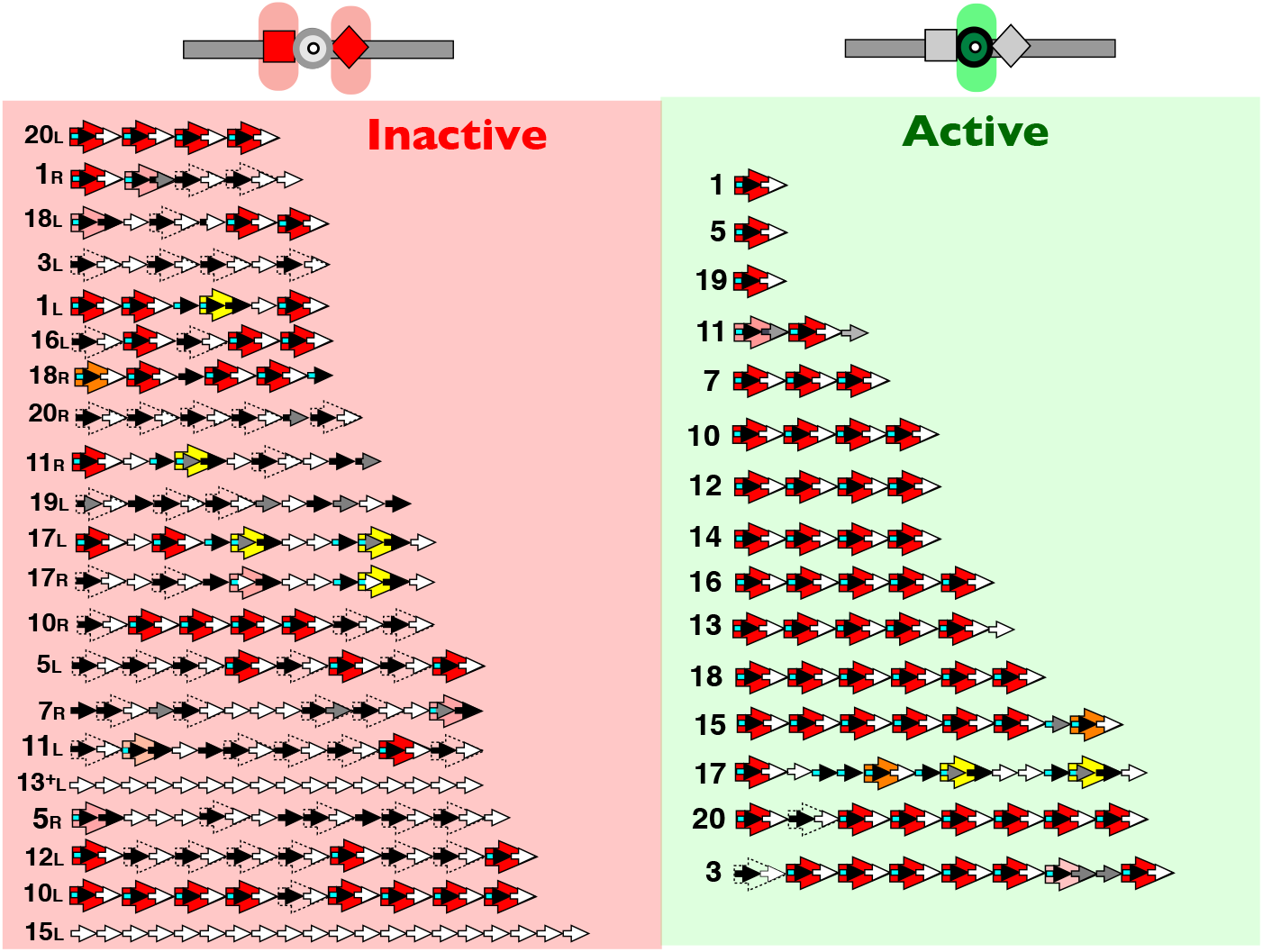
Comparison of the HOR structure of the active HORs and their inactive flanking HORs (immediately to their left or right in the UCSC genome browser [GRCh38]). Active HORs are only included for chromosomes that have flanking HORs. The flanking arrays on chromosomes 13, 14, 21, and 22 are not fully resolved in the UCSC genome browser [GRCh38] (Ziccardi et al. 2016), so a single flanking array (labeled 13^+^) is used for this group of four chromosomes.

**Table 1.** Standing length variation on the human X and Y (Means are from Table 1 and Variances and Coefficients of Variation are calculated from and Figure 4B of Miga et al. 2014). Each sex chromosome has two distinct array cluster groups (groups 1 and 2).

| Chromosome | Array Group | Mean (M) | Variance (V) | Coefficient of Variation (CV) |
| --- | --- | --- | --- | --- |
| X | All | 3.175 | 0.673 | 26.67 |
|  | 1 | 2.698 | 0.183 | 10.84 |
|  | 2 | 4.062 | 0.332 | 21.93 |
| Y | All | 0.803 | 0.204 | 59.22 |
|  | 1 | 0.399 | 0.024 | 44.42 |
|  | 2 | 1.186 | 0.078 | 24.71 |

### Long runs of b-box monomers are absent

Long runs of n-box monomers are found on the Y chromosome (34 in a row, Figure 4) and also on flanking HORs on the autosomes (up to 20 in a row, Figure 9). In the flanking HORs, there are also many long runs of monomers lacking functional b-boxes (including both monomers with broken b-boxes that do not bind CENP-B, and also n-box monomers; Figure 9). In sharp contrast, long runs of b-box monomers are absent in active HORs, with the longest run being a single case of three in a row on chromosome 17 (Figure 4). This avoidance of long runs of b-box monomers (but not monomers lacking them) suggests some form of functional constraint that disfavors these runs. Avoidance of runs of b-box monomers is not predicted by the unequal crossing over model.

### Flanking, inactive HORs are predicted to make weak kinetochores

Monomers containing CENP-B-binding b-boxes are expected to contribute to stronger, more functional centromeres because they increase the level of the kinetochore protein CENP-C recruited to centromeres (Fachinetti et al. 2015). Kinetochore proteins bind DNA at nucleosomes containing the histone H3 variant CENP-A. Fachinetti et al. (2015) showed that CENP-C (a foundational kinetochore protein that recruits down-stream kinetochore proteins; Tanaka et al. 2009) binds to CENP-A containing nucleosomes at two sites: i) the carboxyl end of the CENP-A protein, and ii) near the amino end of this protein where CENP-A, -B, and -C form a triplex in which CENP-C binds DNA-bound CENP-B –but only if CENP-B is bound to the amino end of CENP-A (Supplemental Figure S19). Binding of CENP-C to DNA-bound CENP-B requires a b-box in the DNA linker separating neighboring nucleosomes (Masumoto et al. 1989; Masumoto et al. 1993). When CENP-B is absent, the amount of CENP-C recruited to centromeres is halved because it is absent near the amino end of CENP-A (Fachinetti et al. 2015; Supplemental Figure S19). This reduction in CENP-C is associated with weaker centromeres that have increased miss-segregation rates (Fachinetti et al. 2015). Centromeric repeats with no (or a low density of) b-box monomers are therefore expected to produce weak, poorly functioning centromeres.

Canonical b-box dimers place a b-box adjacent to every nucleosome within an HOR array (see Supplemental Figure S12 in the companion paper; Rice 2019) and this arrangement would feasibly permit maximal recruitment of CENP-C and thereby produce strong centromeres. If currently inactive, flanking HORs were to become activated, the substantially reduced density of monomers containing b-boxes (compared to presently active HORs; Figure 9) would be expected to produce weak, lower-functioning centromeres. This feature indicates that flanking HORs are most feasibly older HORs that have been replaced by currently active, higher functioning HORs. Using the metaphor described earlier, most or all flanking, inactive HORs appear to be graveyards, not nurseries. The difference in predicted centromere strength of active and inactive HORs is not predicted by unequal crossing over model.

## The lifecycle of HORs

The observed structure at active HORs and inactive flanking HORs motivates a hypothesis for a lifecycle during the evolution of centromeric repeats (Figure 10). In this cycle, HORs begin as short dimeric repeats (e.g., the 1-dimer HORs observed on chromosomes 1, 5, and 19) with canonical b/n-box dimeric structure. The high sequence similarity of the dimers that make up the short HORs located on chromosomes 1, 2, 5, and 19 to dimers found on much longer HORs on other chromosomes (Supplemental Figure S18) suggest that new HORs are recruited by ectopic exchange with established HORs, e.g., via ectopic gene conversion or **B**reak-**I**nduced **R**epair (BIR) with template switching. Once established, new, short HORs initially expand only by adding additional b/n-box dimers (not single monomers) with substantial sequence similarity to the existing dimers, i.e., dimers that map to the same cluster in Figure 5 (see also Figure 6 and Supplemental Figure S17). The high sequence similarity between some dimers of long HORs located on different chromosomes (e.g., the X and chromosome 17, and chromosomes 4 and 9; Supplemental Figure S15) suggest that ectopic exchange between chromosomes may contribute to HOR expansion. Once large enough (i.e., > 4 dimers in length), HORs begin to expand by adding both canonical b/n-box dimers as well as unpaired b-box and n-box monomers (with a genetic similarity constraint). At these larger sizes, they also begin to lose functional b-boxes by mutation (i.e., convert canonical b/n-box dimers to broken b/n-box dimers that cannot recruit CENP-B), further reducing canonical b/n-box modular structure. Once there is sufficient proportional loss of canonical b/n-box dimeric structure, these long HORs lose their ability to recruit half of a foundational kinetochore protein CENP-C. As a consequence, these HOR arrays make weaker kinetochores, and are replaced (possibly via centromere drive [Malik and Henikoff 2002] for stronger kinetochores [Iwata-Otsubo et al. 2017]) by short HORs containing canonical b/n-box dimers (originating from pieces of existing, HORs). Once inactive, HOR arrays eventually shrink to complete loss via recurrent deletion pressure (especially due to repair of DSBs via the SSA pathway).

**Figure 10.**
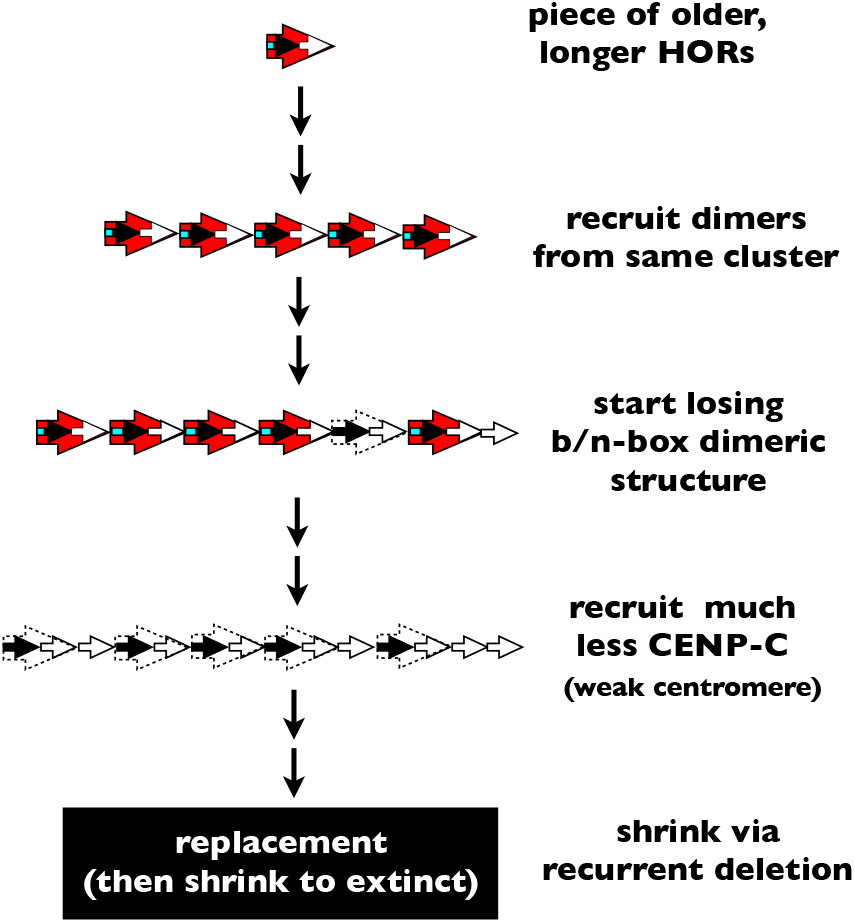
The inferred life history of a human HORs across time.

If the hypothesized sequence of events for a HOR life history is correct (Figure 10), then many questions immediately arise. How and why:

- do HORs grow from short to long?,
- is expansion of short HORs restricted to the addition of b/n-box dimers?
- does canonical b-box/n-box structure decay once HORs become long?
- do new, short canonical b-box/n-box dimers invade and expand within an array of longer, b-box-depleted HORs?
- does the active state (CENP-A epigenetic mark) of an HOR move between new and old HORs?
- do HORs evolve so much more rapidly then the rest of the genome, and sometimes in a punctuated manner?

These questions I address in the companion paper (Rice 2019).

## Evaluating the unequal crossing over model of Smith

### The Smith model is missing an important recombination repair pathway

The foundation of the Smith model for the evolution of repetitive DNA is mitotic, out-of-register recombination between repeat arrays on sister chromatids, i.e., mitotic unequal crossing over (Smith 1976). Unequal crossing over is one outcome of **H**omology **D**irected **R**epair (HDR), which includes repair by the **S**ynthesis-**D**ependent **S**trand **A**nnealing pathway (SDSA; no crossing over), and the **D**ouble **S**trand **B**reak **R**epair pathway (DSBR; repair via the formation of double Holliday structures that sometimes results in crossing over); Fishman-Lobell et al. 1992; Schildkraut et al. 2005. Missing from the Smith model (because it had not yet been discovered) is repair of DSBs via the **S**ingle **S**trand **A**nnealing pathway (SSA; no crossing over). In the long arrays of tandem rDNA repeats in yeast, repair of endonuclease-induced DSBs typically leads to the deletion of one repeat unit per DSB (Ozenberger and Roeder 1991). Subsequent studies support the general conclusion that SSA-repair within many different types of tandem repeats typically leads to small deletions of one repeat unit per DSB (Muchova et al. 2014; Bhargava et al. 2016; Warmerdam et al. 2016). Chakraborty et al. (2016) conclude that SSA is the major DSB repair pathway used within most highly repetitive genomic regions, and that this pathway is favored in these regions because it limits large-scale rearrangements. Although I have not found studies screening for SSA repair of DSBs at human centromeric HORs, there is experimental evidence for SSA repair of DSBs in the centromeric repeats of mice –in combination with repair via HDR (Tsouroula et al. 2016).

The importance of the presence of even low levels of SSA repair at centromeric repeats is that it is expected to act as a deterministic process that continuously erodes the length of HOR arrays –in small steps of one repeat unit per DSB. The unequal crossing over process is length neutral (at the population level) because an unequal crossover lengthens the HOR array on one sister chromatid by the same amount that it shortens the other. When repair of DSBs at centromeric repeats uses both the length-neutral HDR and length-eroding SSA repair pathways, as was found for the centromeric repeats in mice (Tsouroula et al. 2016), HOR arrays are expected to gradually shrink over time. Some factor besides HDR-induced out-of-register sister chromatid recombination is required to explain the evolution and persistence of the multi-megabase-long arrays of centromeric repeats observed at human centromeres (Willard 1991; Miga et al. 2014).

Natural selection for sufficiently long centromeric arrays would be a simple mechanism, to maintain their long length in the face of continuous erosion due to SSA-repair of DSBs. This intuitive solution seems unlikely, however, because the length of most centromeric arrays is far in excess of that needed for normal cellular functioning (Lo et al. 1999; Yang et al. 2000). So some other process is maintaining the extremely large size of human centromeric HOR arrays, despite their continual erosion by SSA repair of DSBs.. In the next section (and more extensively in the companion paper; Rice 2019), I explore the hypothesis that repair of collapsed DNA replication forks via the BIR pathway is this missing process.

### The Smith model cannot account for levels of length polymorphism at the centromeric HOR arrays of the sex chromosomes

The foundation of the Smith model is that most length variation at centromeric HOR arrays arises from unequal crossing over between these arrays on sister chromatids during the repair of DSBs. As described below, a comparison of length variation at the centromeric repeats of the sex chromosomes indicates that an alternative process –repair of collapsed DNA replication forks via the BIR pathway– more plausibly generates this variation.

In general, neutral standing genetic variation increases with an increasing neutral mutation rate (μ, the *gain* parameter for new genetic variants) and an increasing effective population size (Ne, the *sampling error* parameter for segregating genetic variants). Larger Ne decreases the loss rate of extant genetic variants by decreasing sampling error (which causes segregating genetic variants to be lost stochastically across generations). These input and sampling error relationships have been quantified for neutral SNP variation with the result that expected standing genetic diversity (π) is proportional to the product of the gain (μ) and loss (Ne) parameters, i.e., π ~ Ne*μ. (Watterson 1975).

The effective population size (N_e_) of the human Y chromosome (N_e(Y)_) is smaller than that of the X (N_e(X)_) because it is male-limited and hemizygous: causing it to have 67% fewer population-wide copies compared to the X. The effective size is further reduced for the male-limited Y due to: i) lack of recombination along most its length, and ii) stronger sexual selection in males compared to females. These two features both cause elevated levels of Hill-Robertson interfering linkage disequilibrium that further reduces N_e(Y)_ (Wilson Sayres et al. 2014). Collectively, the Y’s male-limited transmission, hemizygosity, and lack of recombination make N_e(Y)_ << N_e(X)_. Empirical studies clearly demonstrate this large inequality in humans based on the much lower standing genetic variation on the Y compared to the X (e.g., Wilson Sayres et al. 2014). This pattern is also seen in other mammals (Hellborg and Ellegren 2004).

The substantially reduced N_e(Y)_ compared to N_e(X)_ causes sampling error to be substantially elevated on the Y compared to the X, leading to the prediction that –when gain parameters (i.e., unequal mitotic crossing over rates between sister chromatids) are similar on both sex chromosomes– the Y will have far less standing variation in centromeric HOR array length compared to the X. A large empirical data set (372 X and 372 Y chromosomes, sampled from around the world; Miga et al. 2014) indicates that the variance (V) in HOR array size is about 70% less on the Y compared to the X (Table 1). But this comparison does not account for the much smaller mean size of the centromeric HOR arrays on the Y chromosome compared to the X (Table 1): and hence its much smaller target size for the DSBs that would generate unequal crossovers. When standing variation is more appropriately measured as the coefficient of variation (CV) to account for array size differences, the Y has as high or higher standing variation compared to the X (means, variances and CVs calculated from Figure 4 and Table 1 of Miga et al. 2014) This surprising and highly counterintuitive finding of high standing variation on the Y compared to the X, despite N_e(Y)_ << N_e(X)_, indicates that the gain parameter for HOR array length variation must be substantially larger on the Y chromosome compared to the X.

The centromeric HOR arrays on the X and Y have similar DNA sequences and chromatin structure. They are: i) composed of AT-rich repeat elements (monomers that are ~170 bp with substantial sequence similarity between the X and Y), ii) complexed with the same kinetochore-associated proteins, and iii) similar in chromatin composition (CENP-A replaces H3 histones at many of their nucleosomes [Scott & Bloom 2014], and the DNA of their repeat arrays is packaged as about one third ‘centrochromatin’, and two thirds heterochromatin; Sullivan & Karpen 2004; Sullivan et al. 2011). All else being equal, this strong similarity in DNA sequence and chromatin structure makes it enigmatic that there could be a large difference in the gain parameter for length variation (unequal crossover rate) in their centromeric arrays.

But all else is not equal. There are two major differences between the centromeric repeats of the X and Y sex chromosomes that could plausibly contribute to large differences in this gain rate. The Y chromosome: i) has an HOR that completely lacks the 17 bp binding sites for CENP-B (b-boxes) that are present at high density on the centromeric repeats of the X and autosomes, and ii) is restricted to males and therefore replicates exclusively within the male germ line. Below I consider two molecular mechanisms by which these differences could combine to feasibly elevate the gain rate for centromeric array length variation on the Y compared to the X. These mechanisms, however, require length variation to be generated by a process other than mitotic unequal crossing over between sister chromatids.

#### Replication-induced DSBs do not contribute to unequal crossing over between sister chromatids

In humans, the male-limited Y differs from the X by experiencing more DNA replications per generation due to substantially more mitoses per generation in the the male vs. female germ line (Wilson Sayers and Markova 2011). Unequal crossing over between sister chromatids is necessarily restricted to the late S and G_2_ phases of the cell cycle when sisters co-occur (note that DSBs are not repaired during M phase of the cell cycle: repair is delayed until G_1_ or the G_1_/S transition stages of the following cell cycle; Bakhoum et al. 2017). This timing constraint would seem to indicate that most DSBs generated during S-phase cannot lead to crossing over between sisters, and therefore that more rounds of DNA replications per generation in the male germ line should not elevate the crossover rate on the Y chromosome. However, DNA replication can potentially induce DSBs that are produced and repaired in late S and G_2_ (when sisters can potentially crossover) due to the intertwining (catenation) of DNA strands from newly synthesized sister chromatids. Post-replication DSBs could be generated due to ligation errors during decatenation of the intertwined sisters by Topoisomerase-II (Liu et al. 2014). However, this route to replication-induced sister chromatid exchange is feasible everywhere in the genome except at centromeres

At human centromeres, intertwined sister chromatid DNA acts as a cohesion agent that – along with cohesin– joins sisters (along centric and/or pericentric regions of chromosomes) until they segregate at anaphase. Correspondingly, and unlike the rest of the genome, the decatenation process at centromeric DNA is delayed until anaphase and early telophase of mitosis (Wang et al. 2010). So despite the intuitive possibility that more DNA replication in the male germ line might elevate sister chromatid crossover rate by generating catenations that later produce DSBs that are repaired during late S and G_2_ phases of the cell cycle, this possibility does not apply to centromeric repeats. This possibility is further reduced by the finding of Chan and Hickson (2011) of no detectable DSBs generated by decatenation of centromeres at the end of mitosis in human cells, as revealed by *γ*H2AX staining. Furthermore, even if some decatination-induced DSBs did occur at centromeres but went undetected by Chan and Hickson (2011), they would be repaired during G_1_ or at the entry of the S-phase of the next cell cycle when sister chromatids are absent: and hence could not generate unequal crossing over between sisters. Therefore, the rate of mitotic crossover should not be substantially influenced by DNA-replication, and hence little affected by the number of mitoses per generation. For this reason, if the Smith model were correct, the smaller effective size of the Y (N_eY_) compared to the X (N_eX_) should lead to much smaller standing variation in HOR array length on the Y: but, as described above, it does not. This finding indicates that some process other than unequal crossing over between sister chromatids (and that operates during DNA replication) is responsible for generating much of the length variation of centromeric HOR arrays.

#### Collapsed replication forks can generate length variation at centromeric repeats

An important molecular process generating DSBs during DNA replication is fork-stalling followed by fork-collapse (fork-stalling/collapse; Syeda et al. 2014; Cannan and Pederson 2016). Unlike the two-ended DSBs generated by factors like free radicals or topoisomerase-II ligation errors, fork-collapse during DNA replication generates one-ended DSBs that are repaired by **B**reak-**I**nduced **R**epair (BIR; Costantino, et al. 2014). Repair of these one-ended DSBs generates length variation in tandem repeat arrays via out-of-register reinitiation of DNA replication (Kobayashi 2014). BIR repair of collapsed replication forks has been studied extensively in the rDNA repeats of budding yeast and found to be biased toward array expansion rather than contraction (Kobayashi 2014). Empirical evidence indicates that fork-stalling/collapse is common during the replication of centromeric DNA in humans (Crosetto et al. 2013; Aze et al. 2016) and nonhumans (Greenfeder and Newlon 1992; Mitra et al. 2014), as described in more detail in the companion paper (Rice 2019). BIR repair of collapsed DNA replication forks would feasibly have two results: i) it generates length variation that increases with more mitoses per generation (as occurs on the Y compared to the X), and ii) it could feasibly counterbalance array shrinkage due to SSA repair of DSBs due to its bias toward array expansion over contraction (as indicated at yeast rDNA arrays; Kobayashi 2014).

Although the average number of cell divisions per generation is more that an order of magnitude higher in human males compared to females (Wilson Sayres and Makova 2011), this asymmetry alone is probably insufficient to fully explain the paradox of the Y centromeric arrays displaying near parity in their level of length variation compared to the X, despite N_e(Y)_ << N_e(X)_. This conclusion comes from the fact that SNP variation on these same chromosomes is much lower on the Y than the X (Wilson Sayres et al. 2014), despite DNA replication contributing substantially to the total SNP mutation rate (Cui et al. 2012) and the same mitoses/generation difference between the sex chromosomes applies to SNPs. However, the absence of CENP-B binding sites within the centromeric array of the human Y (but not the X) could feasibly magnify the effect of the Y’s higher cell divisions per generation.

#### CENP-B can influence the rate of collapsed replication forks

CENP-B almost certainly has function(s) outside its role in kinetochore formation because the protein is found in most mammal species (Yoda et al. 1992; Casola et al. 2008) yet its binding site is absent at the centromeres of most of these species (Haaf et al. 1995; Alkan et al. 2011; Kugou et al. 2016). CENP-B m ed i ates su rvei l l an ce for retrotransposons in fission yeast (Cam et al. 2008) and it may have a similar function in mammals (Kipling and Warburton 1997; Casola et al. 2008). In fission yeast, CENP-B binds to AT-rich regions of LTR transposons and recruits other proteins that reduce the fork-stalling/collapse that is required for the transposons to replicate (Zaratiegui et al. 2011). Assuming CENP-B produces a similar phenotype in humans, its absence on the centromeric repeat array on the Y, and presence on the X, would raise the relative rate of fork-stalling/collapse on the the Y compared to the X (on a per bp basis), and this in turn would elevate the input rate of length variation. Together, the Y chromosome’s i) higher cell divisions per generation, and ii) feasibly higher rate of fork-stalling/collapse due to lack of CENP-B binding, could plausibly combine to contribute substantially to the unexpectedly high variation in HOR array length found on the low-Ne Y chromosome –but as described above, these DNA replication features would not be expected to increase the length variation input rate via conventional mitotic unequal crossing over between sister chromatids.

In sum, the high HOR array length variation on the Y relative to the X, despite N_e(Y)_ << N_e(X)_, is enigmatic under the Smith model that assumes unequal crossing over between sister chromatids is responsible for most length variation at centromeric repeats. This finding, however, is plausible when one instead assumes that fork-collapse during DNA replication (followed by out-of-register BIR) is responsible for this length variation, especially if CENP-B bound to centromeric DNA reduces this fork-collapse phenotype, as it does in fission yeast.

### The Smith model cannot account for high between-species sequence divergence observed at the X-linked centromeric HOR arrays of humans and chimps

In most cases, gradual sequence divergence via base pair substitutions and small indels cannot be measured between the orthologous HORs of humans and chimps because almost all centromeric HOR arrays have experienced a replacement by distantly related HOR sequences within one or both lineages (Archidiacono et al. 1995). The X chromosome is the sole exception. As described in more detail in the companion paper, X-linked HOR sequences have diverged at a rate that is at least 5-fold faster than the rest of the euchromatic and heterochromatic genome. This highly elevated divergence rate is not predicted by Smith’s unequal crossing over model –unless centromeric regions have: i) exceptionally high mitotic crossover rates between sister chromatids (for which I have found no evidence), and ii) high CpG content (Arbeithubera et al. 2015; which is not the case for human monomers (Supplementary Table S1). The high base pair substitution rate is predicted, however, if most length variation at human HORs is generated by fork-collapse during DNA replication followed by re-initiation of DNA replication via out-of-register BIR. This elevation occurs because BIR generates long replication tracts with error-prone DNA polymerases that increase the base substitution mutation rate over 1,000-fold compared to DNA replication without fork-collapse (Sakofsky et al. 2012). I will explore this possibility in the companion paper (Rice 2019).

### Smith model does not predict the many levels of centromeric HOR structure

The patterns observed when comparing the consensus sequences from all of the active HORs to each other, and to their flanking, inactive HORs (consensus sequences from from GRCh38), can be compared to what would be expected under the Smith (1976) model of evolution within a tandem repeat (that is based on out-of-register crossover between sister chromatids). This neutral model predicts that the centromeric repeats will be composed of essentially random sequences with no substantive structural constraint except the avoidance of harmful sequences, such as those producing fold-back loops that stall replication forks. In sharp contrast, and as described in the preceding sections, I found the consensus sequences of active centromeric repeats to be highly structured, at multiple levels (Figures 7 and 9), despite sequence divergence among monomers as large as 35% (Figure 5). This diversity of structure is summarized in Box 1.

#### Box 1. Structure at different levels within the consensus sequences of all centromeric HORs.

1. All consensus monomers within all active centromeric HORs have the same head-to-tail orientation, i.e., there are no inversions among the monomers that make up centromeric HORs. All monomers also contiguously align to a single consensus sequence, indicating that there are no large inversions within individual monomers. The persistence of these structural patterns despite strong sequence divergence among monomers suggests functional constraint.
2. The spacing among b-box elements (including broken b-boxes) is highly nonrandom: they are separated by an integer number of monomers (never two b-boxes within one monomer) and a spacing of every other monomer strongly predominates (Figure 2). This nonrandom spacing of b-boxes within HORs supports the hypothesis that their placement is functionally constrained.
3. Monomers that do not contain a b-box at their 5’ end have a different conserved sequence that is 19 bp in length (n-box, Supplemental Figure S10B). These conserved n-boxes support the hypothesis of functional constraint on the 5’ end of monomers lacking b-boxes.
4. There is highly significant length differentiation between n-box and b-box monomers: b-box monomers are significantly shorter (averaging ~2 bp shorter) and 8.6 times more variable in length than n-box monomers. Differences in mean and variance of the length of n-box and b-box monomers, despite substantial sequence divergence among them, support the hypothesis that they have different functional constraints.
5. Runs of b-box monomers of at least moderate length (> 3 in a row) are absent (longest run is a single case of 3 in-a-row on chromosome 17). In sharp contrast, exceptionally long runs of n-box monomers are present on the Y chromosome (34 n-box monomers in-a-row; Figure 4) and on the flanking arrays of the autosomes (up 20 n-box monomers in a row; Figure 9). In addition, long runs of monomers that lack a functional b-box (runs of no-b-box monomers that are a mixture of n-box monomers and those with broken b-boxes) are common in the flanking, inactive HORs (Figure 9). The absence of long runs of b-box monomers (but not no-b-box monomers) supports the hypothesis of functional constraint on the spacing of b-box and n-box monomers within HORs.
6. Short HORs (≤ 10 monomers in length) have a nearly perfect modular structure composed of canonical b/n-box dimers, whereas longer HORs usually deviate from this modularity by having reduced density of functional b-box monomers (i.e., including broken b/n-box dimers and/or lone monomers) (Figure 7). This pattern supports the hypothesis of different functional constraint on short vs. long HORs.
7. The HOR on the male-limited Y chromosome is the only one of the active HORs that lacks a high density of b-boxes (Figure 4). This pattern of b-boxes present within the centromeric repeats of the X and all of the autosomes but absent on the Y chromosome is also seen in the laboratory mouse (*Mus musculus domesticus;* Pertile et al. 2009). The absence of b-boxes on only the Y chromosome among distantly related taxa is unlikely by chance alone and indicates that monomer sequences are not random, i.e., they are influenced by some aspect of Y-linkage.
8. The consistent position of both n-box and b-box elements near the 5’ end of monomers (Supplemental Figure S10) positions them within the linker DNA regions between nucleosomes (Tanaka et al. 2005; Hasson et al. 2013; Fujita et al. 2015; Henikoff et al. 2015). The fact that the position of these short sequences has not drifted away from this linker position, despite substantial sequence divergence among monomers (and when b-boxes are absent –as they are on the HOR on the Y chromosome), supports the hypothesis that monomer sequence is nonrandom and somehow promotes this consistent nucleosome positioning.
9. It is common for repeat elements (monomers or dimers) to be most closely related to repeat elements from a different HOR on another chromosome rather than those from the same HOR on the same chromosome (Supplemental Figures S15, S16), suggesting that they frequently originate by ectopic exchange (e.g., gene conversion or Break-Induced Repair with template-switching; Hastings 2010) rather than unequal crossing within an HOR. Nontrivial levels of ectopic exchange are also indicated by several chromosome groups ([1, 5 and 19]. [13 and 21], and [14 and 22]) that each sharing a common HOR sequence. Also, the dimers of the shortest active HORs have strong sequence similarity to b/n-box dimers from longer active HORs on different chromosomes (Supplemental Figure S18). These patterns support the hypothesis that ectopic exchange of HOR parts among chromosomes, rather than unequal crossing over within an HOR, contributes substantially to the evolution of HOR sequences.
10. All active HORs on the X and autosomes have a high density of b-box monomers and there is experimental evidence for a strong association between b-box presence and centromere strength: HORs lacking b-boxes, like the one on the Y chromosome, or those with b-boxes but experiencing experimentally lowered levels of cellular CENP-B, make weak centromeres with elevated mis-segregation rates (Supplemental Figure 19; Fachinetti et al. 2015). This pattern supports the hypothesis that HOR sequence is nonrandom and can strongly influence kinetochore structure and function.
11. In comparison to active HORs, inactive flanking HORs are on average (Figure 9): 1) longer, 2) lower in the density of functional b-boxes, 3) higher in the density of n-box monomers (with strongly conserved n-boxes), 4) reduced in modular b/n-box structure, and 5) predicted to form weak kinetochores with high mis-segregation rates if they were to become active (Fachinetti et al. 2015). These structural differences between active and inactive HORs support the hypothesis that there are functional differences between their sequences.
12. While most HORs are a monotonous repetition of the same sequence of monomers, some (like the HOR shared by chromosomes 1,5, and 19) are highly heterogeneous and contain subregions of expanded, longer HORs. The dimers in these regions of longer HORs are closely related to dimers from longer HORs on other chromosomes. These observations support the hypothesis that some HORs are currently in the process of increasing their length by recruiting new dimers via ectopic exchanges between chromosomes –rather than out-of-register unequal crossing over within an HOR.
13. Active HOR arrays are far longer than required for normal cellular functioning (Lo et al. 1999; Yang et al. 2000). This extreme size is not predicted by the unequal crossing over model, especially with the expected perpetual contraction of HOR arrays due to SSA repair of DSBs. Adding to the Smith model selection for sufficient array size would not be expected to generate the extreme sizes observed in nature.

Collectively, the diverse structural patterns listed in Box 1 are not predicted by the Smith model of random sequence evolution solely via unequal exchange between sister chromatids, and indicate substantial functional constraints on HOR sequence and organization.

### Nucleosome positioning over the highly diverged repeat sequences of the Y centromere indicate non-random sequence evolution

The 34 monomers that make up the centromeric HOR on the Y chromosome all lack b-boxes and differ in sequence from each other by as much 28% (Supplementary Table S1). Despite this substantial sequence divergence, Hasson et al. (2013) found highly conserved nucleosome positioning over orthologous nucleotide positions among the Y’s monomers. This conserved nucleosome positioning, despite considerable sequence divergence, would not be expected if the divergence were unconstrained and randomly generated by unequal crossing over, as predicted by the Smith model.

## Conclusions

Many attributes of the human centromeric HORs appear to be in conflict with the widely accepted Smith (1976) model for the evolution of centromeric tandem repeats via unequal crossing over between sister chromatids. First, the Smith model predicts an essentially random sequence (excepting intrinsically harmful sequences, such as those that are self-complementary) for the monomers that make up human centromeric HORs, yet as described here, they have substantial non-random structure at many organizational levels: i) within monomers (e.g., strongly conserved b- and n-boxes located near the 5’ end of monomers that form the linker regions between nucleosomes), ii) between monomer types (e.g., differing means and variance in the length of b- and n-box monomers, suggesting they have different constraints on their length), iii) between pairs of adjacent monomers (e.g., b/n-box dimers are the predominate HOR substructure), iv) among tandem groups of monomers (e.g., all monomers within HORs have the same head-to-tail orientation), v) among HORs of different length (e.g., levels of b/n-box dimeric structure of HORs are strongly correlated with HOR length), and vi) between active and inactive HORs (e.g., inactive HORs are on average longer, have more broken b-boxes, and form shorter arrays [usually much shorter]). Additional evidence for functional constraint on centromeric HORs is i) the absence of runs of >3 monomers with b-boxes, ii) the strong level of nucleosome positioning on b/n-box dimers as well as on the 34 monomers of the Y chromosome despite substantial sequence divergence among these dimers and Y-linked monomers, iii) the observation that active centromeres are predicted to be stronger (recruit more kinetochore proteins) than inactive, flanking HORs. The Smith model also does not include the SSA pathway for the repair of DSBs, and when this process operates during at least some DSB repairs, centromeric repeats are predicted to continually shrink –yet this perpetual contraction has not been reported. One might rescue the underlying logic of the Smith model by simply adding constraints on features like array length, b-box position within monomers and density within HORs, and so on. However, the foundation of the Smith model is unequal crossing over between sister chromatids, which is assumed to generate the substantial length variation we see among extant centromeric repeat arrays. But the high length variation on the Y compared the the X despite N_e(Y)_ << N_e(X)_ is incompatible with the assumption that most length variation at centromeric repeats is generated by unequal crossing over between sister chromatids. The much larger-than-needed size (for cellular functioning) of centromeric repeats is also inconsistent with Smith’s length-neutral unequal crossing over model and indicates that some deterministic lengthening process is continually expanding HOR arrays and offsetting their continual shrinkage via SSA repair of DSBs. Alternatively, these patterns are consistent with a different array expansion/contraction mechanism that is biased toward array expansion: out-of-register repair of collapsed DNA replication forks via the BIR pathway during the replication of tandem repeats. For all of these reasons it seems appropriate to consider alternatives to the Smith model for the evolution of human centromeric HORs. In the companion paper (Rice 2019), I consider such an alternative that is based on length variation that is generated via BIR repair of collapsed DNA replication forks.

## Acknowledgements

I thank two undergraduates working in my laboratory (Roselyn Wu and San Ha Lee) for assistance during the exploratory stages of the sequence analysis. I received helpful comments of the early ideas that motivated this paper when I presented a seminar on centromere evolution at the Fred Hutchinson Cancer Research Center, especially from Harmit Malik. Additional comments were provided by Steve Henikoff on an early draft of the manuscript and by Urban Friberg and Andrew Stewart on a near-final draft of the manuscript. Copy-editing assistance was provided by Kathryn Schoenrock.

---

**Supplemental Figure S1.**
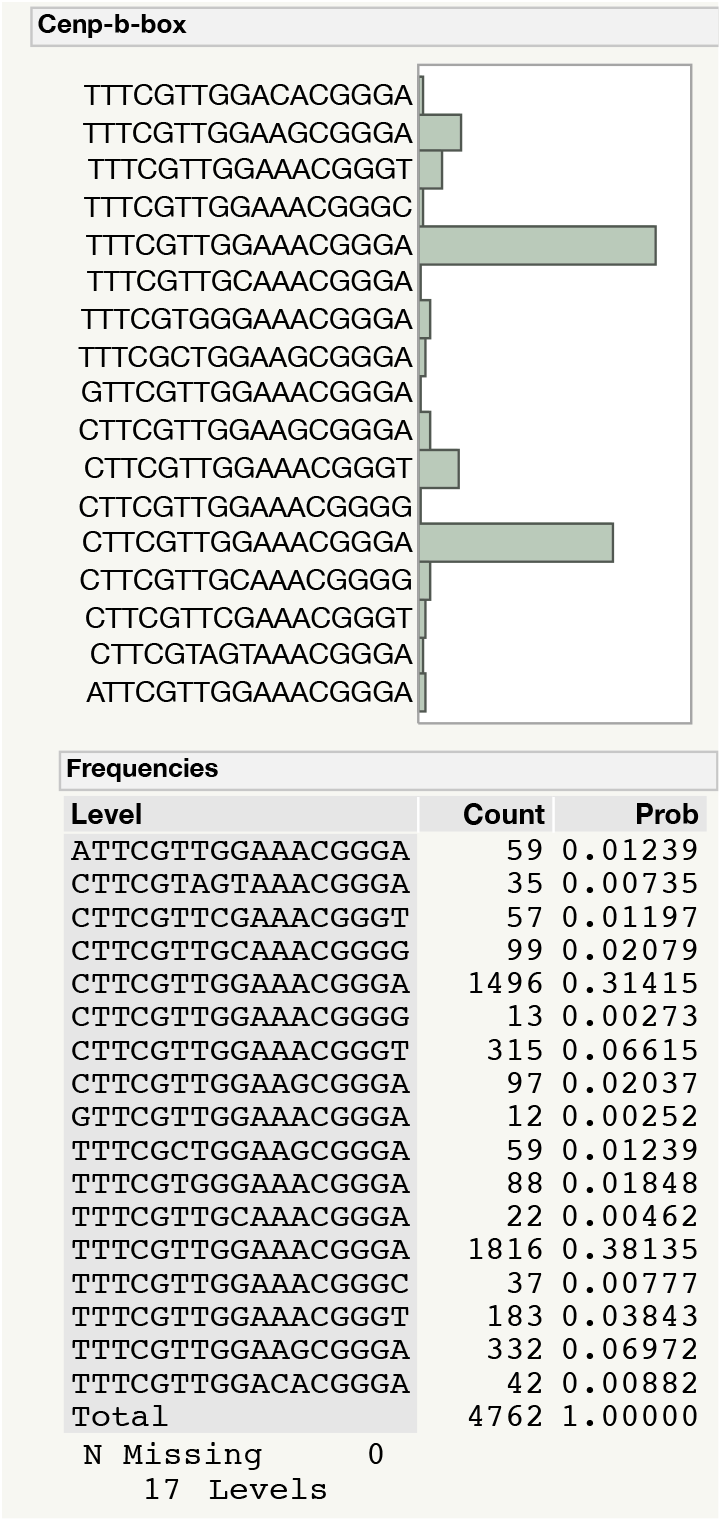
The most commonly found CENP-B binding b-box sequences (consensus = (5’-Y**TTCG**TTGG**A**AU**CGGG**A-3’) from a sample of Illumina reads from a homozygous human female hydatidiform mole (SRR1514950, Chaisson et al. 2015). In vitro studies indicate that only the properly spaced boldface bases in this sequence are required to allow binding of the CENP-B protein (Masumoto et al. 1989; Masumoto et al. 1993). Based on this minimal binding sequence, I first searched a large sample of Illumina 101 bp reads (8.76 x 10^7^) for all 17 bp hits containing the properly spaced 9 bp sequence needed for binding of the CENP-B protein. Only 17 functional b-box sequences were common in the hydatidiform mole genome, and only two of these predominated. These 17 CENP-B-box sequences were used for the BLAST analysis of long PacBio SMRT reads from the same hydatidiform mole.

**Supplemental Figure S2.**
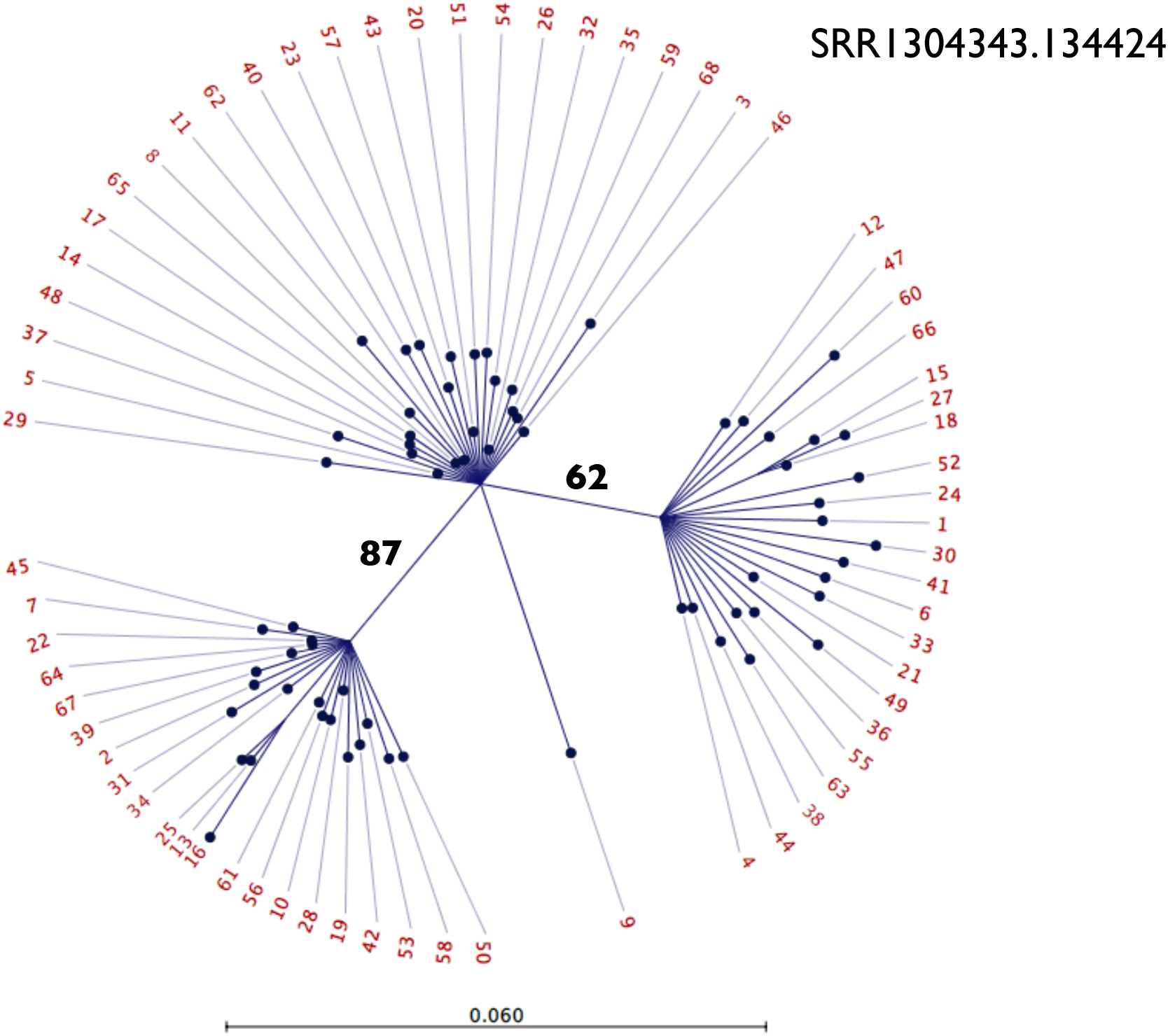
Clustering (neighbor-joining tree) of 68 sub-reads produced by cutting a single PacBio read at all BLAST hit sites for the 17 bp b-box sequence (using the 17 predominant sequences shown in Supplemental Figure S1). All branches with less than 60% bootstrap support were collapsed and bootstrap support for remaining clusters are shown. BLAST of the consensus of each of the three clusters (on GRCh38 at the NCBI web site https://blast.ncbi.nlm.nih.gov/Blast.cgi?PAGE_TYPE=BlastSearch&BLAST_SPEC=OGP__9606__9558&LINK_LOC=blasthome) mapped to the centromeric region of chromosome 7.

**Supplemental Figure S3.**
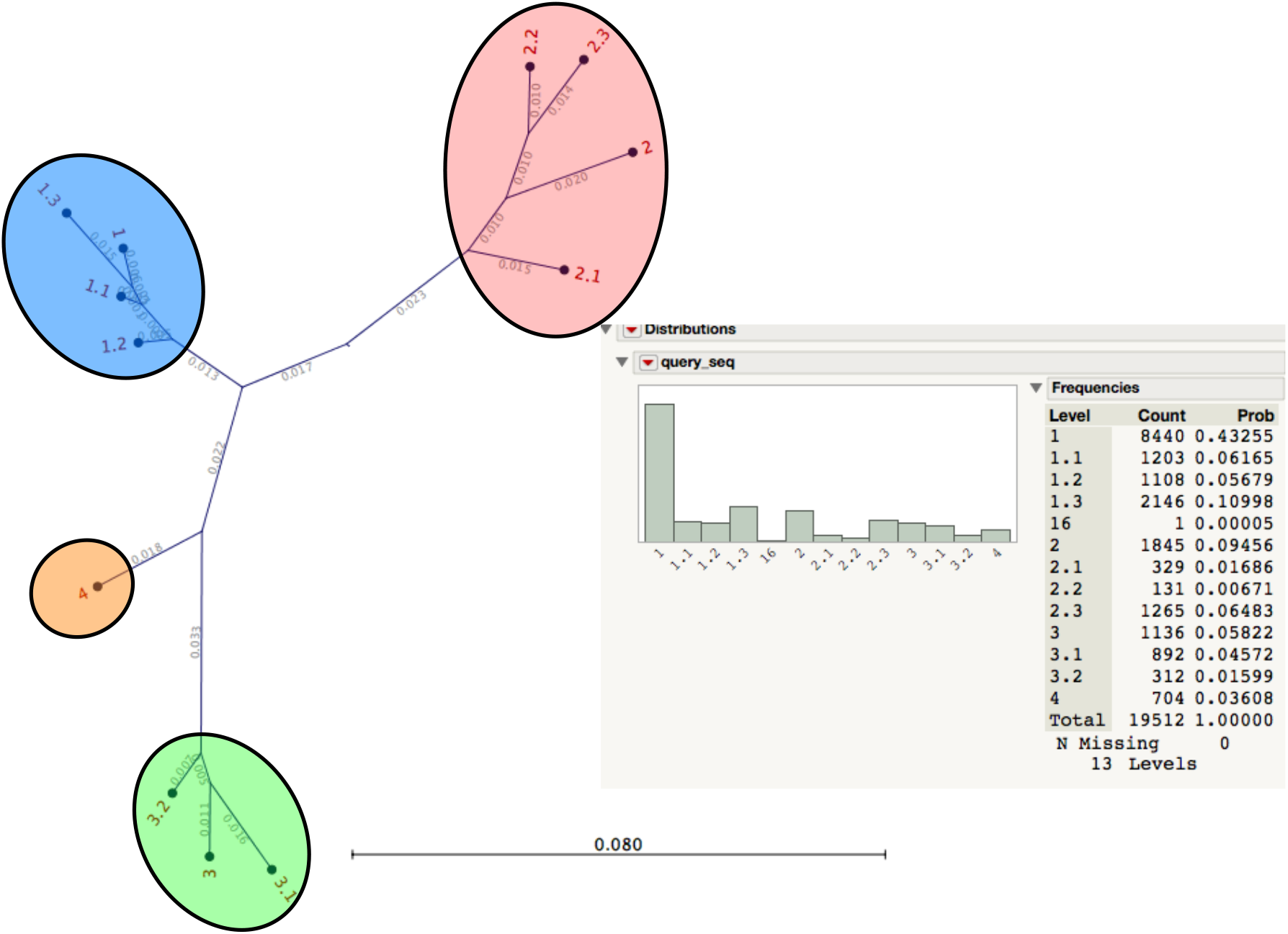
In a sample of 510 PacBio reads containing a total of 19.5 x 10^3^ dimers that mapped to chromosome 1, I found four major sequence clusters of b-box/no-b-box dimers (labeled 1 through 4 with numbers after the decimal point indicating less common subgroups). Most (about two thirds) of the dimers mapped to the blue cluster of 1 - 1.3).

**Supplementary Figure S4.**
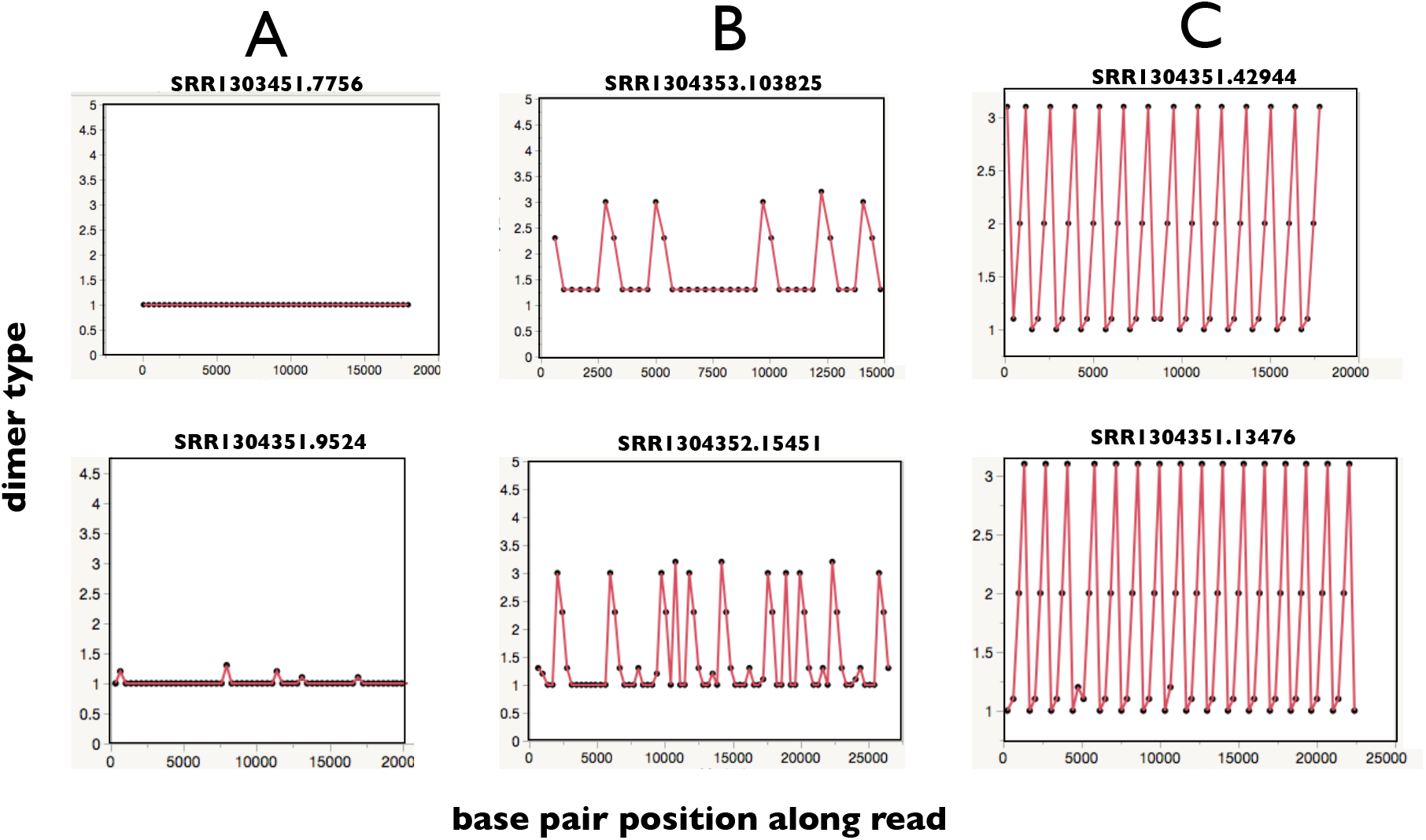
Representative repeat graphs (two for each type) from PacBio reads containing centromeric sequences mapping to chromosome 1 (that also map to chromosomes 5 and 19). **A.** About a quarter of the PacBio reads were homogeneous repeats of the same dimer or dimers from the same cluster with nearly the same sequence (see clusters in Supplemental Figure S3). **B.** However, most reads were a disorganized mixture of dimers that mapped to two or more clusters (see Supplemental Figure S3). **C.** About 2.5% of the reads were a highly homogeneous 4-dimer HORs with two dimers from clusters one and one each from clusters two and three.

**Supplementary Figure S5.**
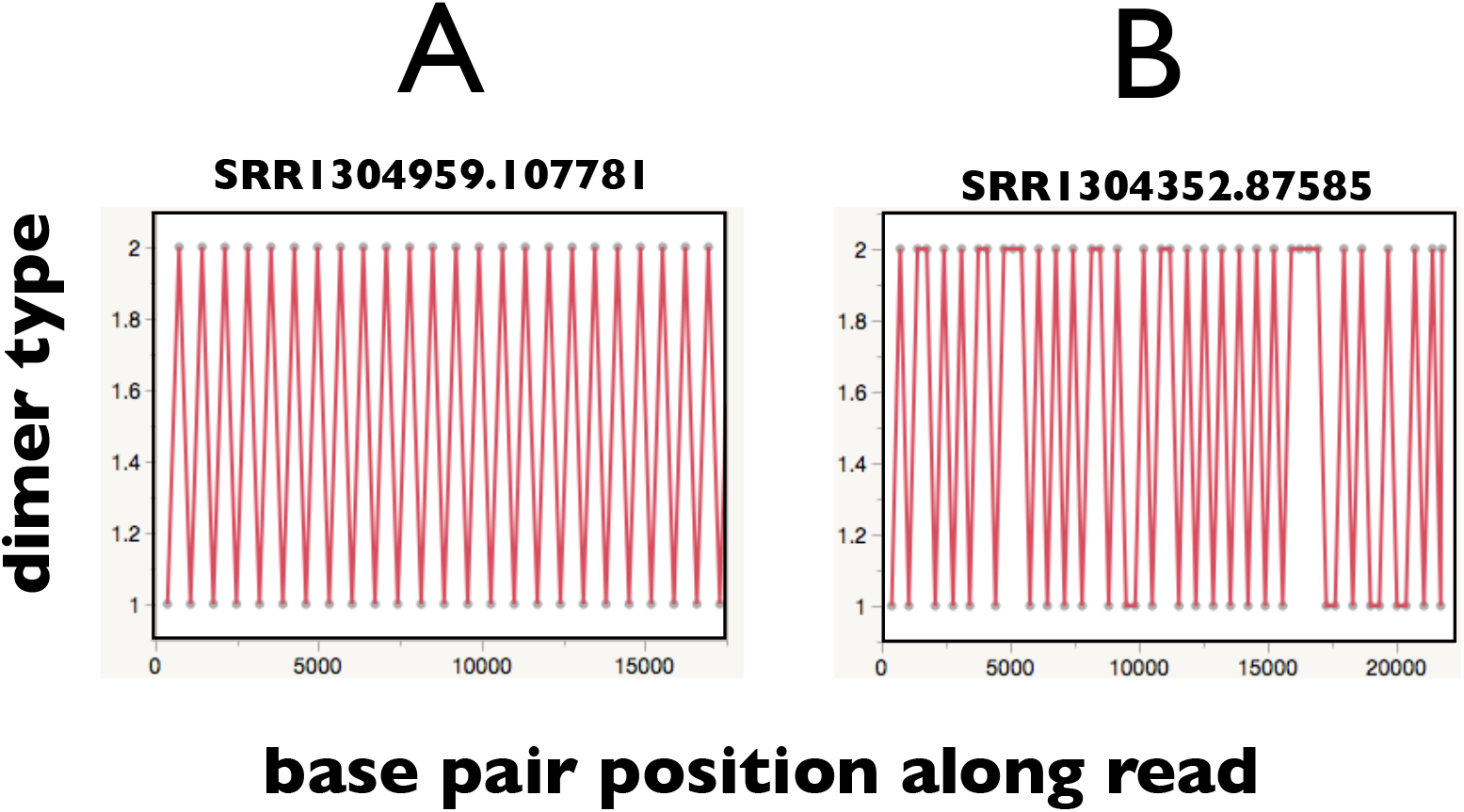
Representative repeat graphs from PacBio reads containing centromeric sequences mapping to chromosome 2. **A.** Many PacBio reads were long homogeneous stretches of the consensus two-dimer HOR reported in Supplementary Table S1 (dimer 2_a = 1 and dimer 2_b = 2). **B.** But most reads were heterogeneous with frequent, short tandem repeats of the same dimer interspersed with more common two-dimer HOR.

**Supplementary Figure S6.**
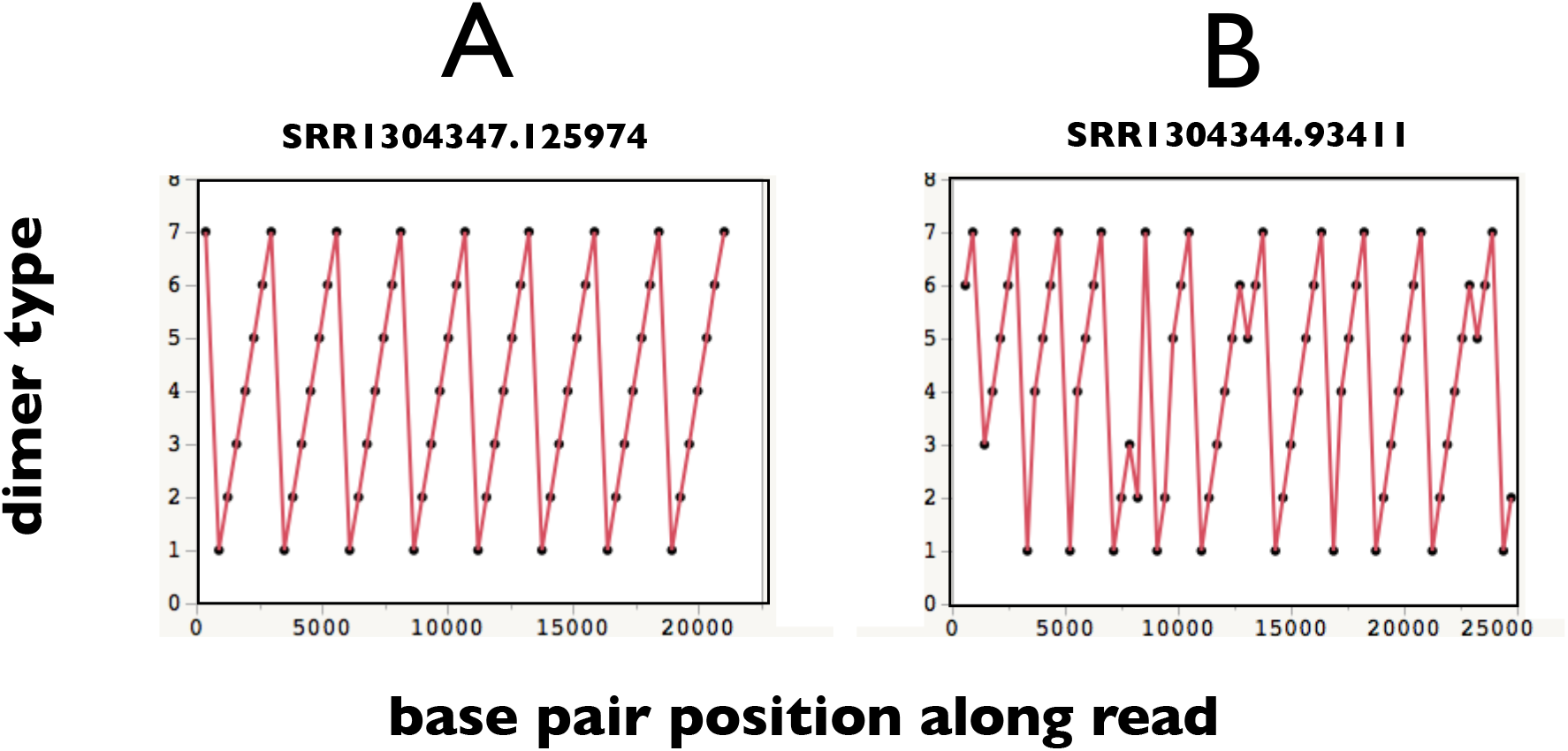
Representative repeat graphs from PacBio reads containing centromeric sequences mapping to chromosome 9. In the graphs, using the labeling from Supplemental Table 1, dimer 9_a = 1, 9_b = 2, 9_c = 3, etc., but “dimer 9_f” includes the 6^th^ dimer plus a following lone monomer. **A.** Many PacBio reads were long homogeneous stretches of the consensus seven-dimer HOR reported in Supplementary Table S1. **B.** But most reads were heterogeneous with frequent indels of one or more dimers.

**Supplemental Figure S7.**
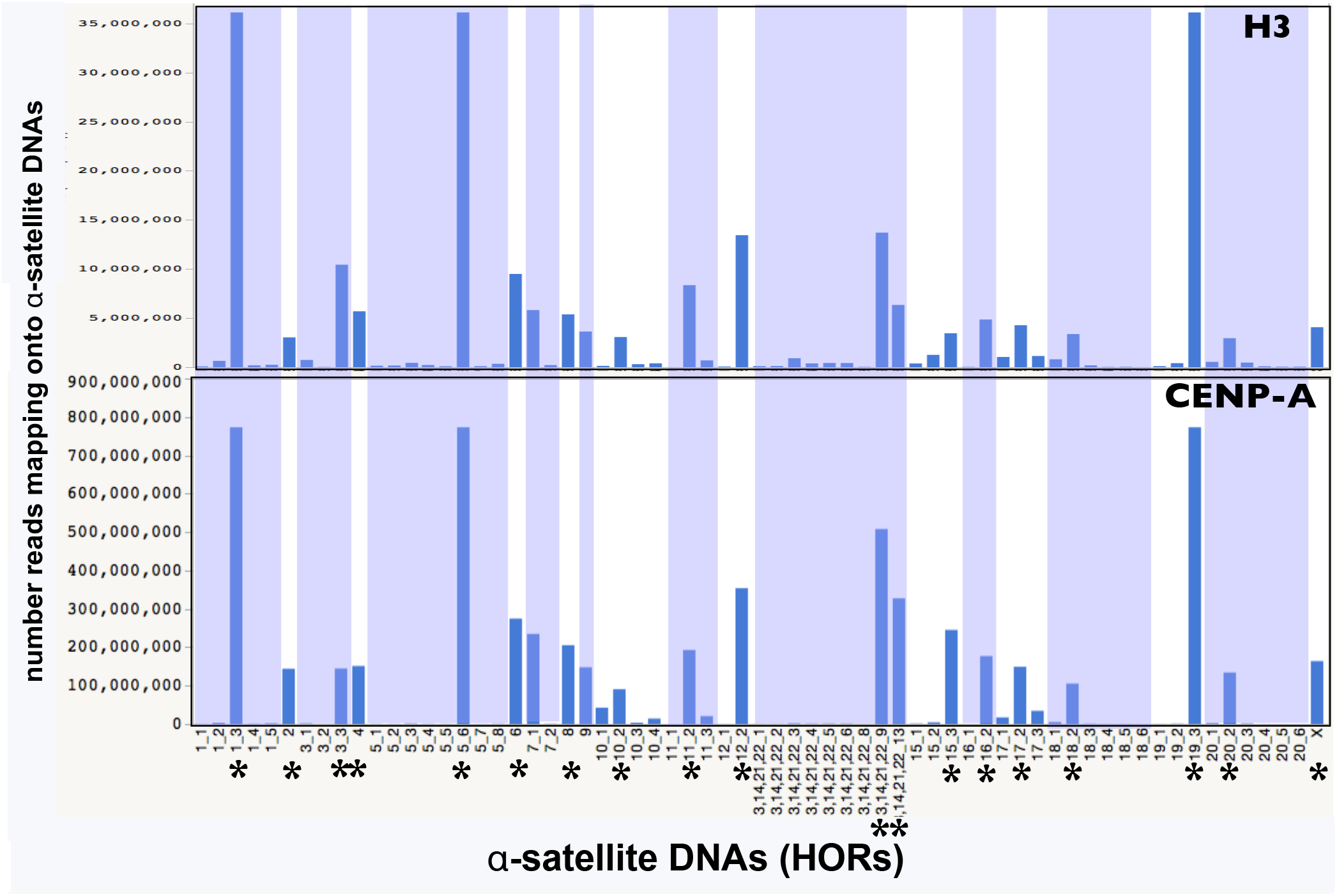
ChIP-seq data compiled from Supplementary Table 4 of Nechemia-Arbely et al. (2017). The DNA sample was MNase digested to mononucleosomes prior to sequencing. The top (H3) sample are Illumina sequences (merged 100 bp paired-end reads) precipitated with antibodies against histone H3 and the bottom (CENP-A) sample was precipitated with antibodies against CENP-A. Numbers along the X axis (i.e., A_B) denote HOR array positions: ‘A’ denotes chromosome and ‘B’ denotes the centromeric HOR array’s position (1 = 1^st^, 2 = 2^nd^, …) in the left-to-right order seen on the UCSC genome browser [GRCh38]. The Y axis is the number of reads mapping to different HORs (read depth). The asterisks denote the region containing the consensus HORs identified here from the PacBio reads of the hydatidiform mole genome (CHM1, Supplementary Table S1). Variation in bar heights (read depths) in the top panel (H3) depict variation in the relative sizes of HORs, but noting that those for chromosomes (1, 5, and 19) and (13,14, 21, and 22) are higher because they combine reads from the multiple chromosomes within each group. Note that the UCSC genome browser does not resolve HORs on chromosomes (13 and 21) and (14 and 22) as separate groups. Bar heights in the lower panel measure read depth in the CENP-A-precipitated DNA, which is expected to be substantial only at active centromeric repeats. In general, each chromosome has one large HOR (strong peaks in top, H3 panel) and this HOR is the only functioning centromeric sequence (strong peaks in bottom, CENP-A panel). Multiple smaller peaks seen on chromosomes 10, 11, and 17 in the bottom panel (CENP-A) are commonly associated with high sequence similarity to the larger, active HOR and may be false positives –but they they may represent joint use of more than one HOR array. The two large peaks seen on the chromosome (13,14,21,22) group represents two active HORs: one on chromosomes 13 and 21 (13_14_21_22_9) and the other on chromosomes 14 and 22 (13_14_21_22_13) (Ziccardi et al. 2016).

**Supplemental Figure S8.**
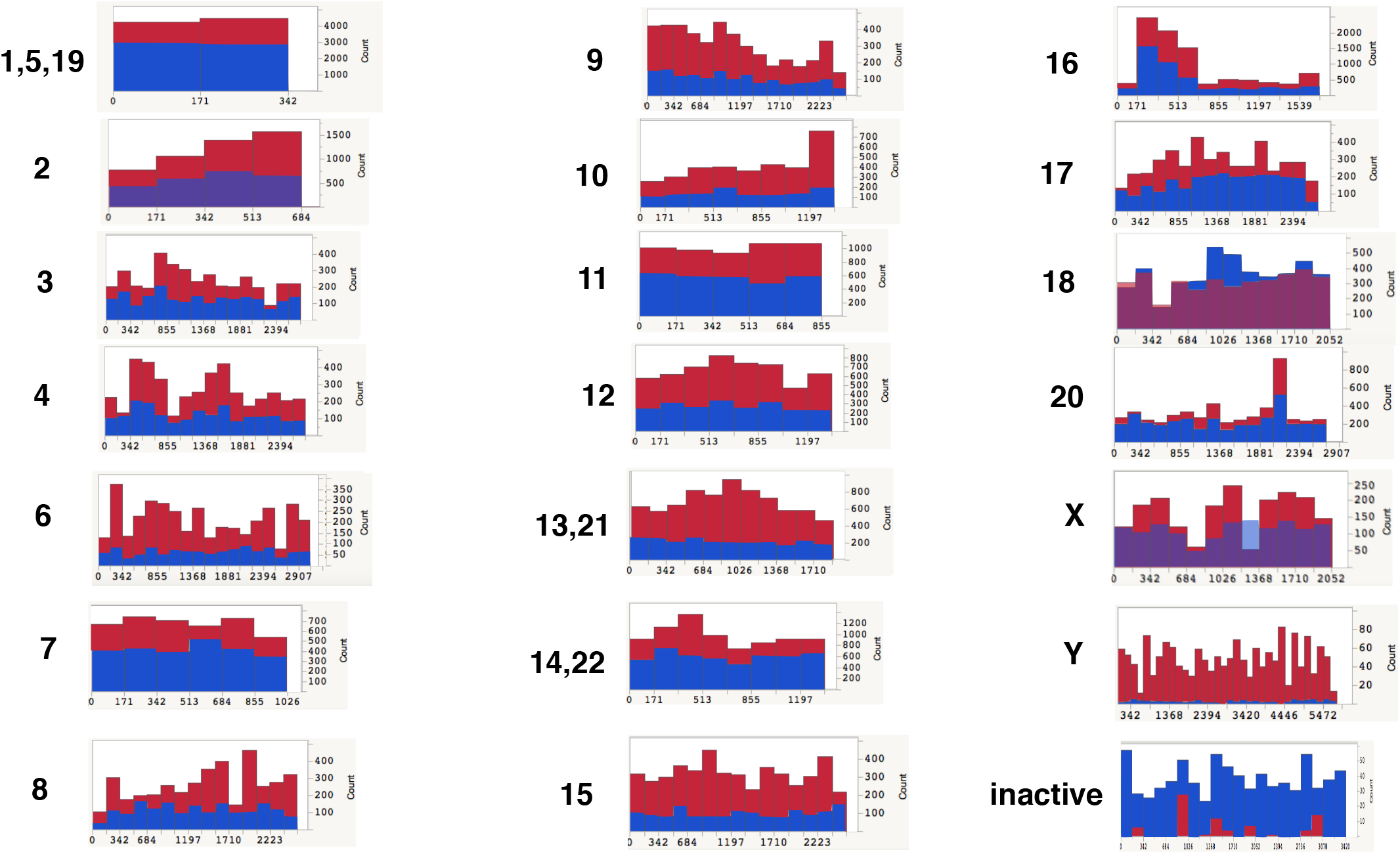
SRA files from a ChIP-seq study by Henikoff et al. 2015 were used to look for CENP-A enrichment at the HORs listed in Supplemental Table 1. DNA was MNase digested to mononucleosomes prior to sequencing. A total of 4.76 x 10^6^ Illumina 100 x 100 bp paired-end reads from either the ChIP or Input SRA files were blasted against each of 20 large-N HORs identified in Supplemental Table 1. Blast hits needed to be at least 97 bp long and match at a level of at least 97% to be scored. Red = ChIP sample and Blue = Input sample. Because each HOR array has only about 400 CENP-A containing nucleosomes, the ChIP/Seq ratio is reduced at larger HOR arrays, i.e., CENP-A signal is diluted in larger arrays (compare the tiny HOR array on chromosome Y to all other, larger HOR arrays). Variation in read depth among monomers of the same HOR is due in part to sequence divergence between the molar genome and the HuRef lymphoblastoid line. Large peaks in chromosomes 16 and 20 are due to strong sequence similarity of these monomers to the HORs on chromosomes (1,5,19) and 2, respectively. All monomers of all HORs show a strong CENP-A enrichment compared to control, indicating that they are active centromeric sequences. The inactive control is the HOR immediately to the left of the active HOR array on chromosome 15, as shown in the UCSC genome browser (GJ212851.1 [GRCh38]) and Supplemental Figure S7.

**Supplemental Figure S9.**
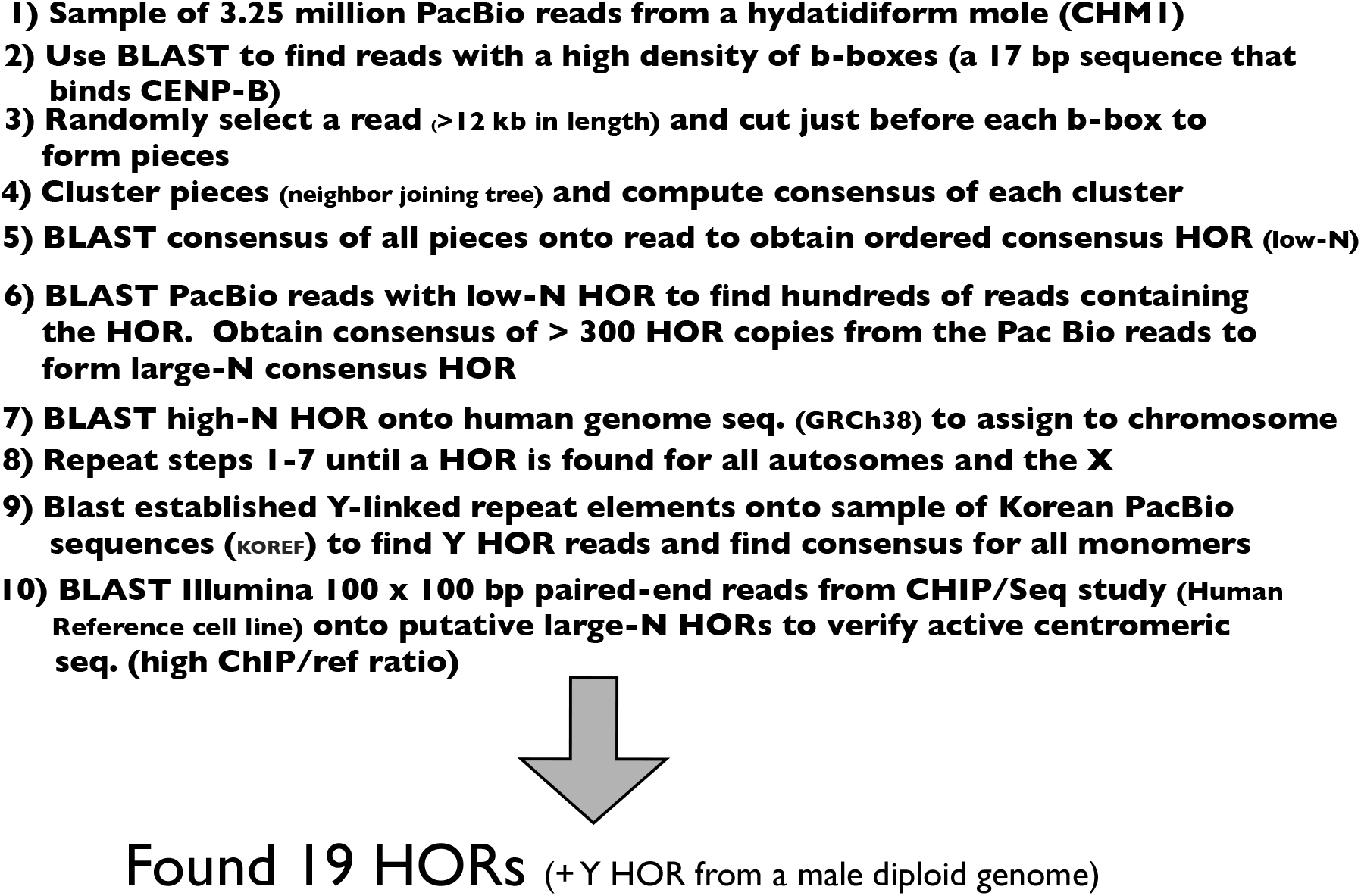
Summary of the pipeline used to identify consensus sequence of all of the active HORs within a single homozygous human genome (and the Y chromosome from a diploid male) from long PacBio reads.

**Supplemental Figure S10.**
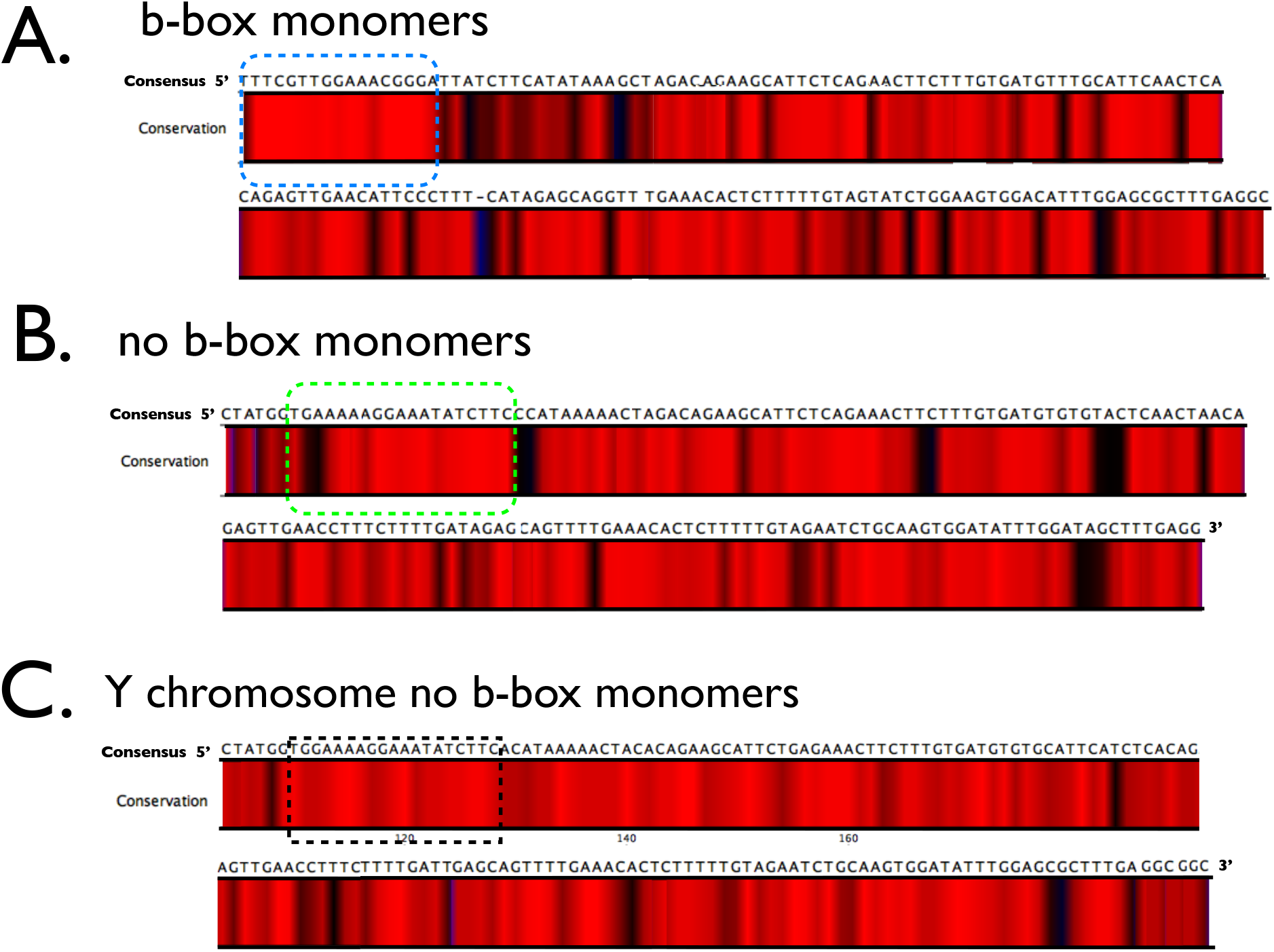
Heat maps of sequence conservation of all b-box **(A)** and no-b-box **(B)** monomers from canonical b/n-dimers with b-boxes that bind CENP-B, i.e., the b-box satisfies -TTCG----A--CGGG-. The 17 bp b-box is strongly conserved at all positions except the first, which as predicted by the consensus sequence, was virtually always C or T. I also observed a strongly conserved 19 bp “n-box” sequence (T[G/A][G/A]AAAAGGAAATATCTTC) displaced 6 bp downstream compared to the b-box position. **C.** The 19 bp “n-box” sequence is also conserved within the 34 no-b-box monomers that make up the consensus sequence of the active HOR on the Y chromosome.

**Supplemental Figure S11.**
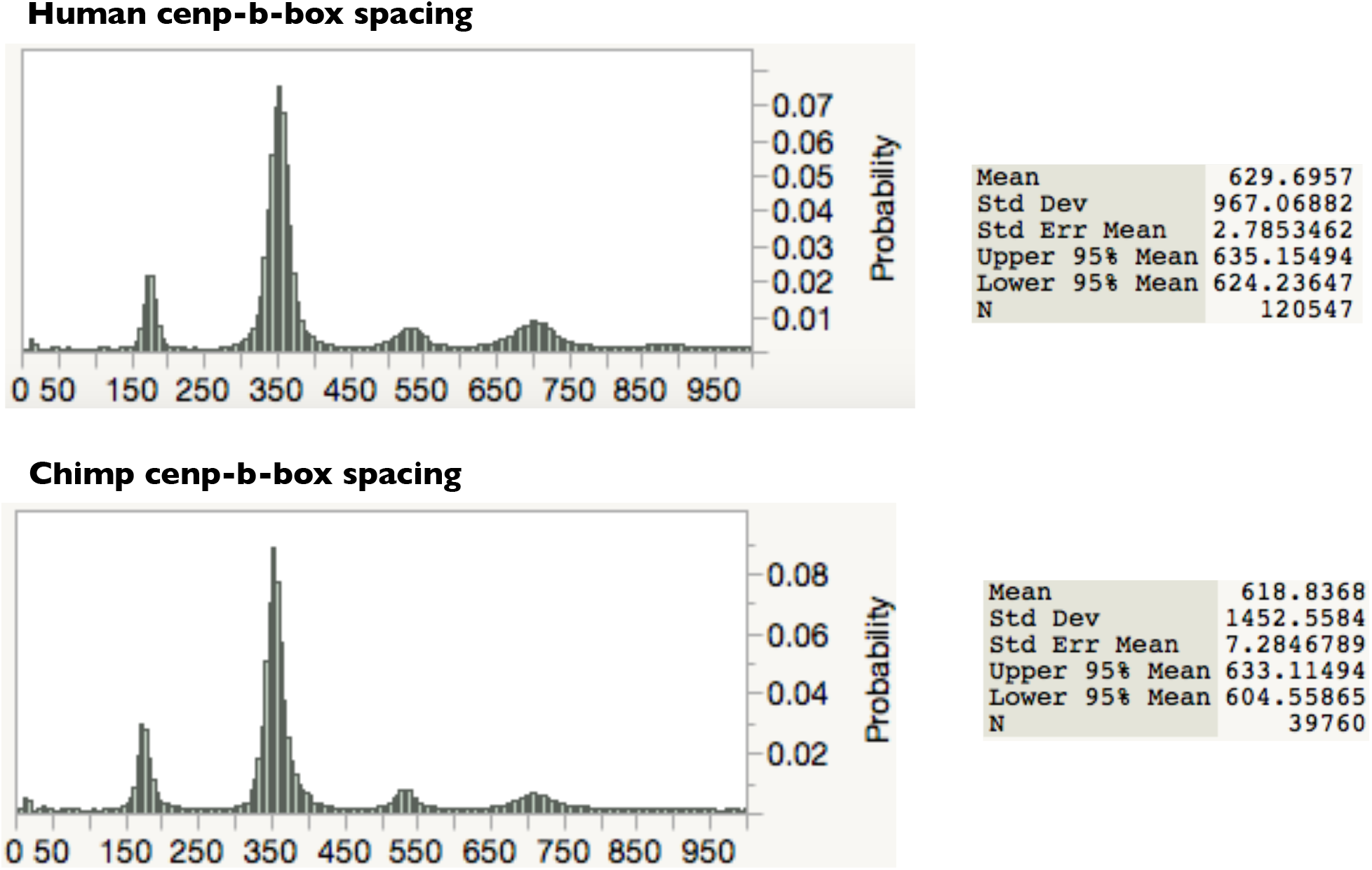
The distribution of the spacing between b-boxes (in a sample of 120,547 b-boxes located on PacBio reads >15kb in length [SRR13043]) in humans is highly similar to that seen in chimps ((in a sample of 39760 b-boxes located on PacBio reads >15kb in length [SRR5269456]). Note that small insertion errors are common in PacBio reads, causing the expected modes (at multiples of ~170 bp) to be slightly larger.

**Supplemental Figure S12.**
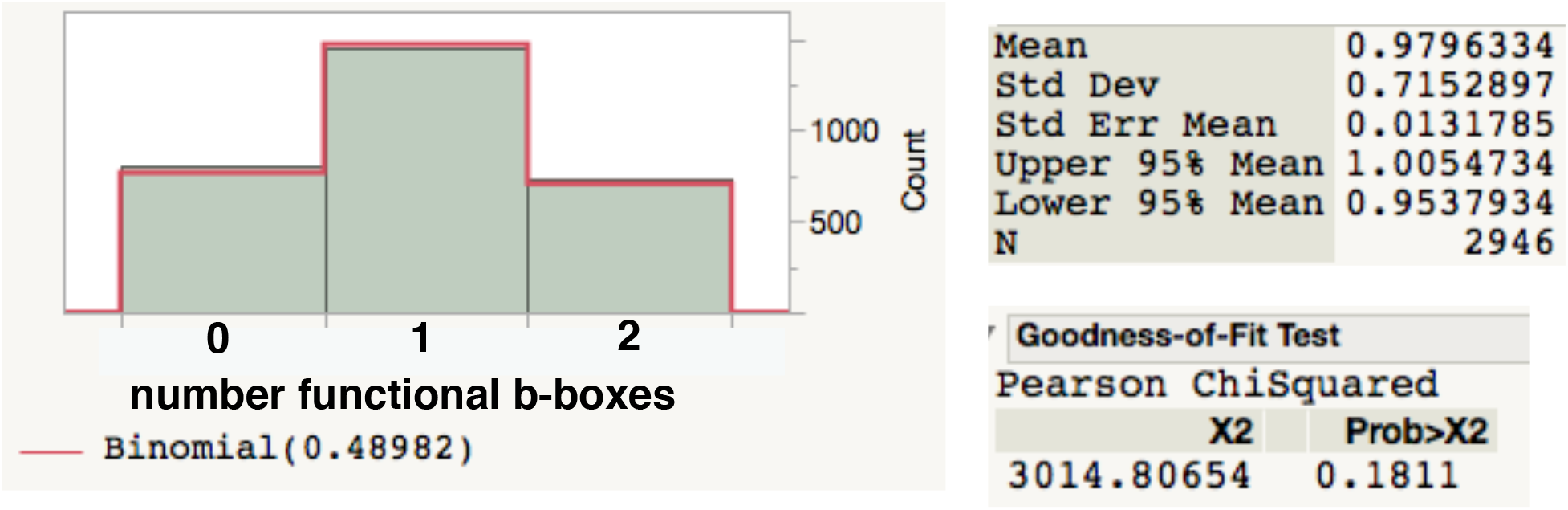
The distribution of the number of functional b-boxes (i.e., those predicted to bind CENP-B because they contain the minimal binding sequence (-TTCG----A--CGGG-) found in a sample of 2946 side-by-side mouse monomers (137 bp sequences: 120 bp monomers starting at the 17 bp b-box [functional or non-functional] followed by the first 17 bp of the adjacent monomer) found in 150 bp shotgun sequence reads (illumina reads located in NCBI file SRR7015105). The red line is the random expectation (based on a binomial distribution). Note that about half of the b-boxes are predicted to be non non-binding for CENP-B, and there is no significant deviation from the null random distribution. Assuming that monomers are CENP-B-binding with probability 0.9796, and they are randomly placed along an centromeric repeat array, then 76.6% of monomers will be included in a b-box/no-b-box dimer or no-b-box/b-box dimer.

**Supplemental Figure S13.**
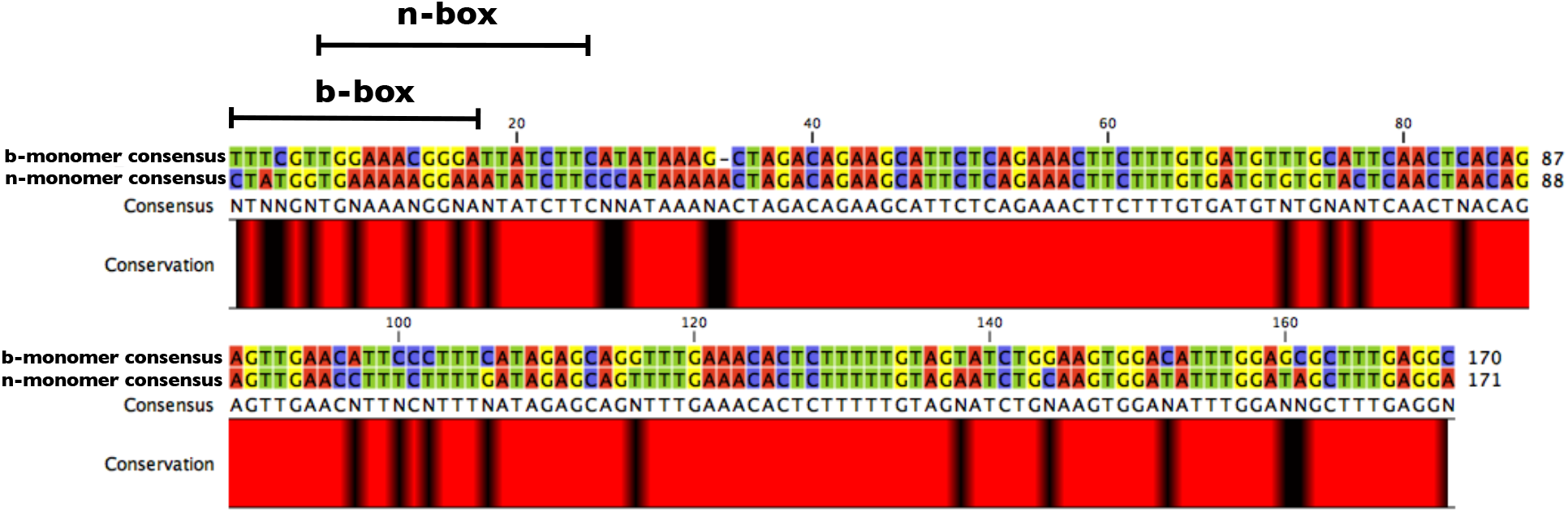
The consensus b-box and n-box monomers (from all canonical b/n-dimers in Supplemental Table S1) have diverged by 15.8% and much of this divergence is outside the b-box/n-box region.

**Supplemental Figure S14.**
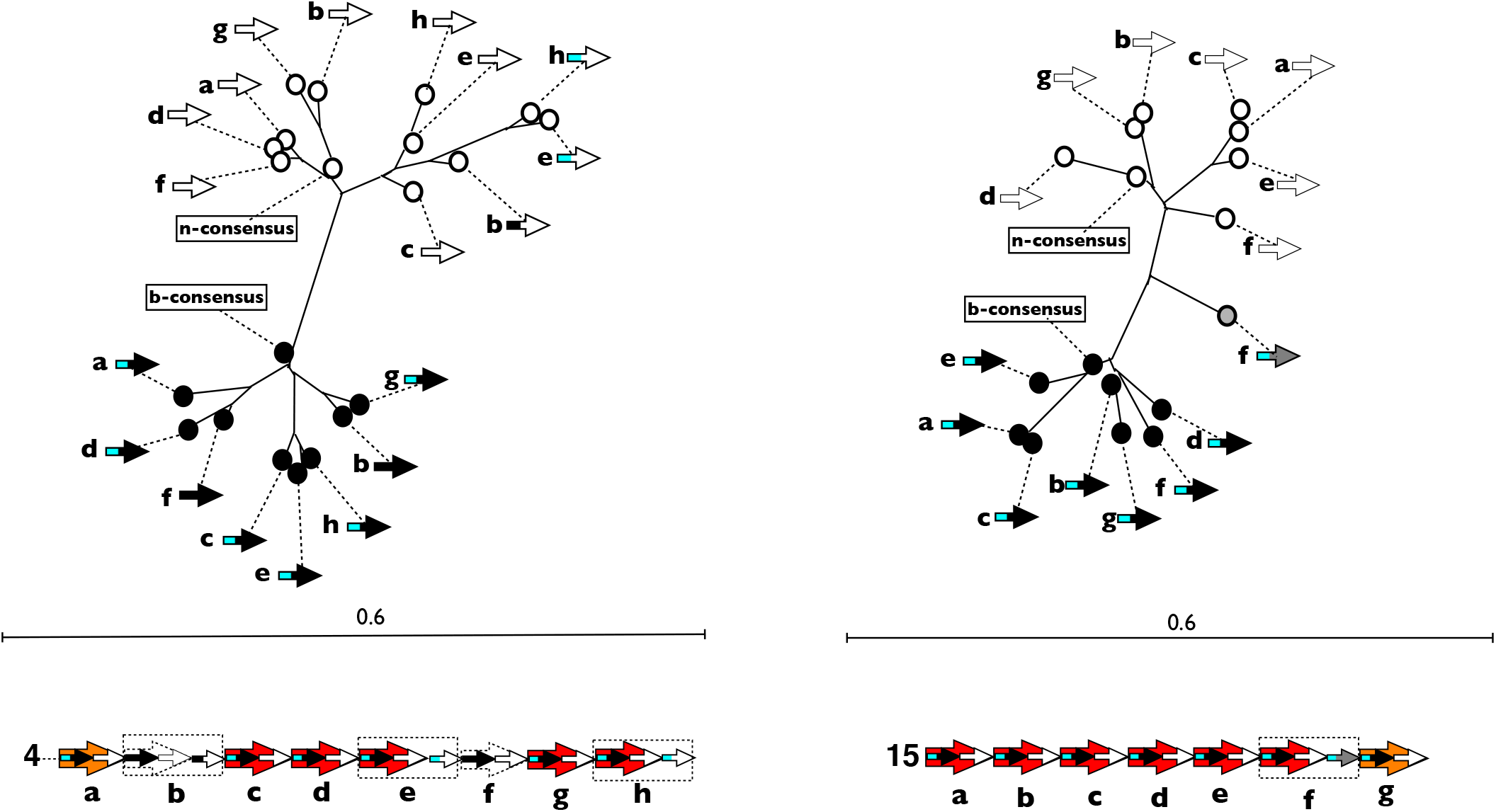
Examples of the cluster classification of monomers (chromosomes 4 [left] and 15 [right]). Each diagram is a neighbor joining tree of all monomers in the large-N consensus HOR. Also included is the consensus b-box monomer and the consensus n-box monomer from the corresponding dimer cluster of Figure 5. Black-filled circles depict b-monomers, white-filled depict n-monomers, and grey are intermediate. Arrow symbols as defined in Figure 3.

**Supplemental Figure S15.**
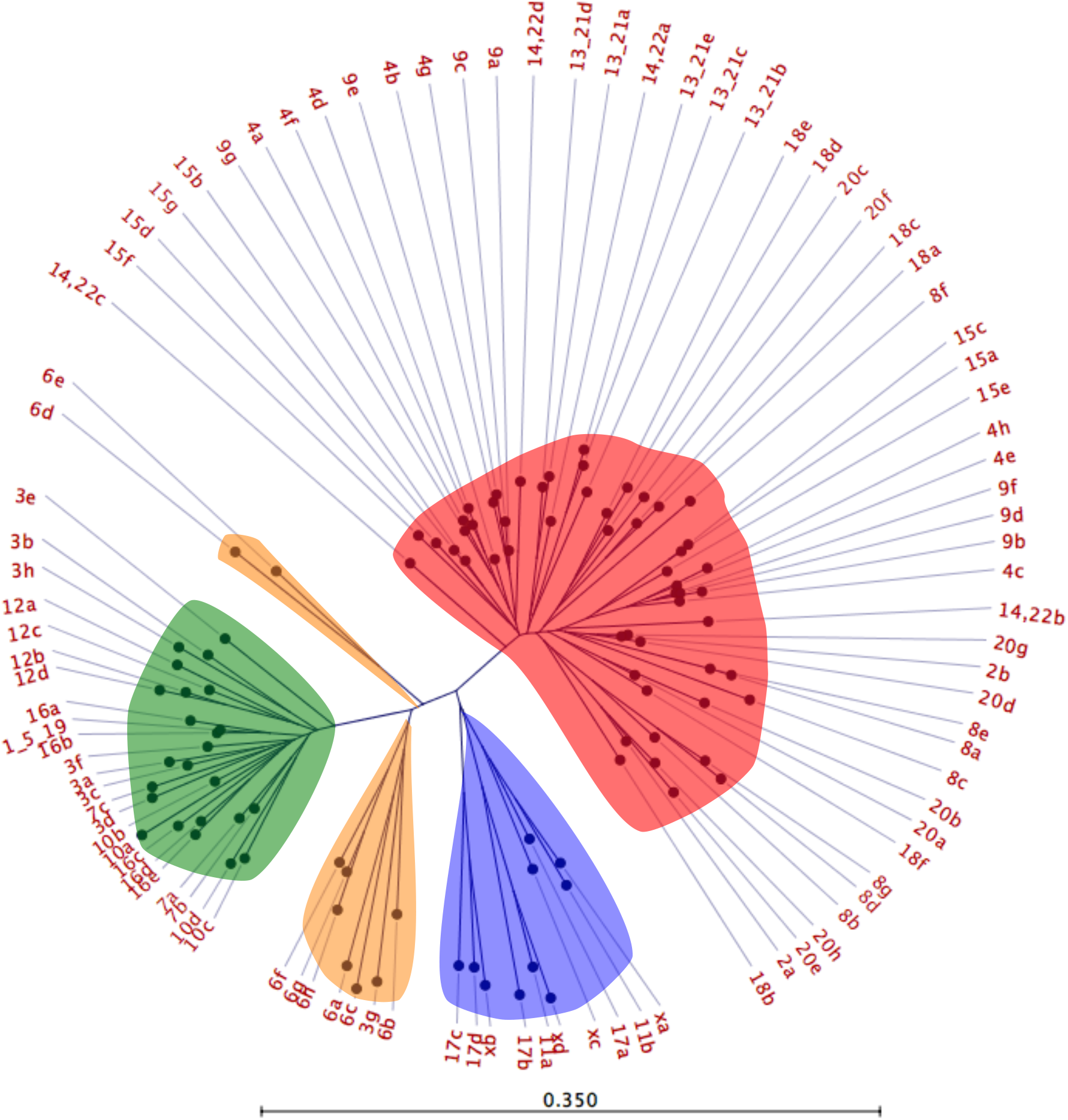
The same figure as Figure 5 in the main article but with individual dimers identified. Dimers are labeled a, b, c, …, where dimer a is the first from the left in Figure 4, b is second from left, and so on.

**Supplemental Figure S16.**
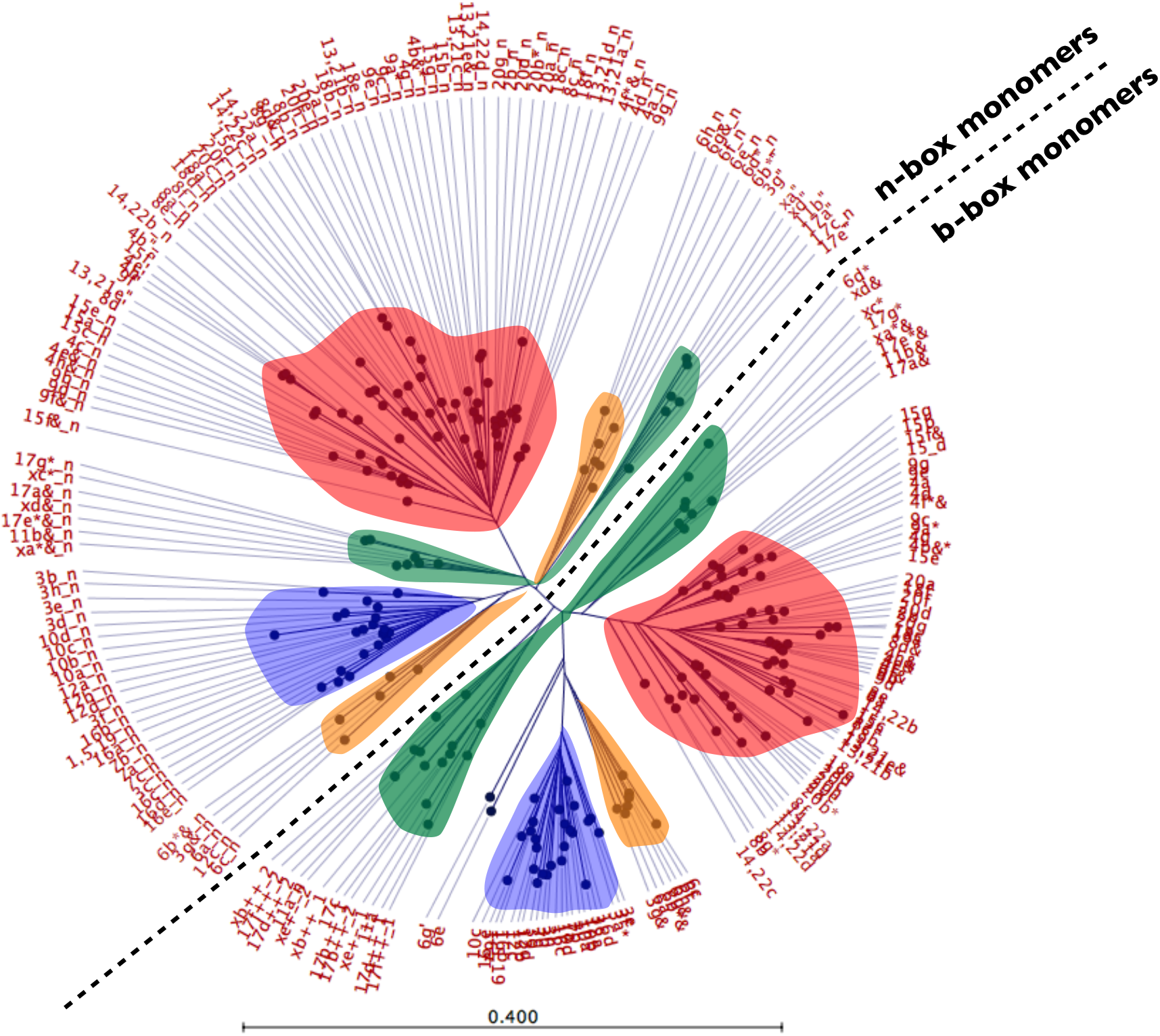
A neighbor-joining tree of all monomers. Clustering of monomers closely matches that of the dimers shown in Figure 5. Monomer names as in Supplemental Table 1.

**Supplemental Figure S17.**
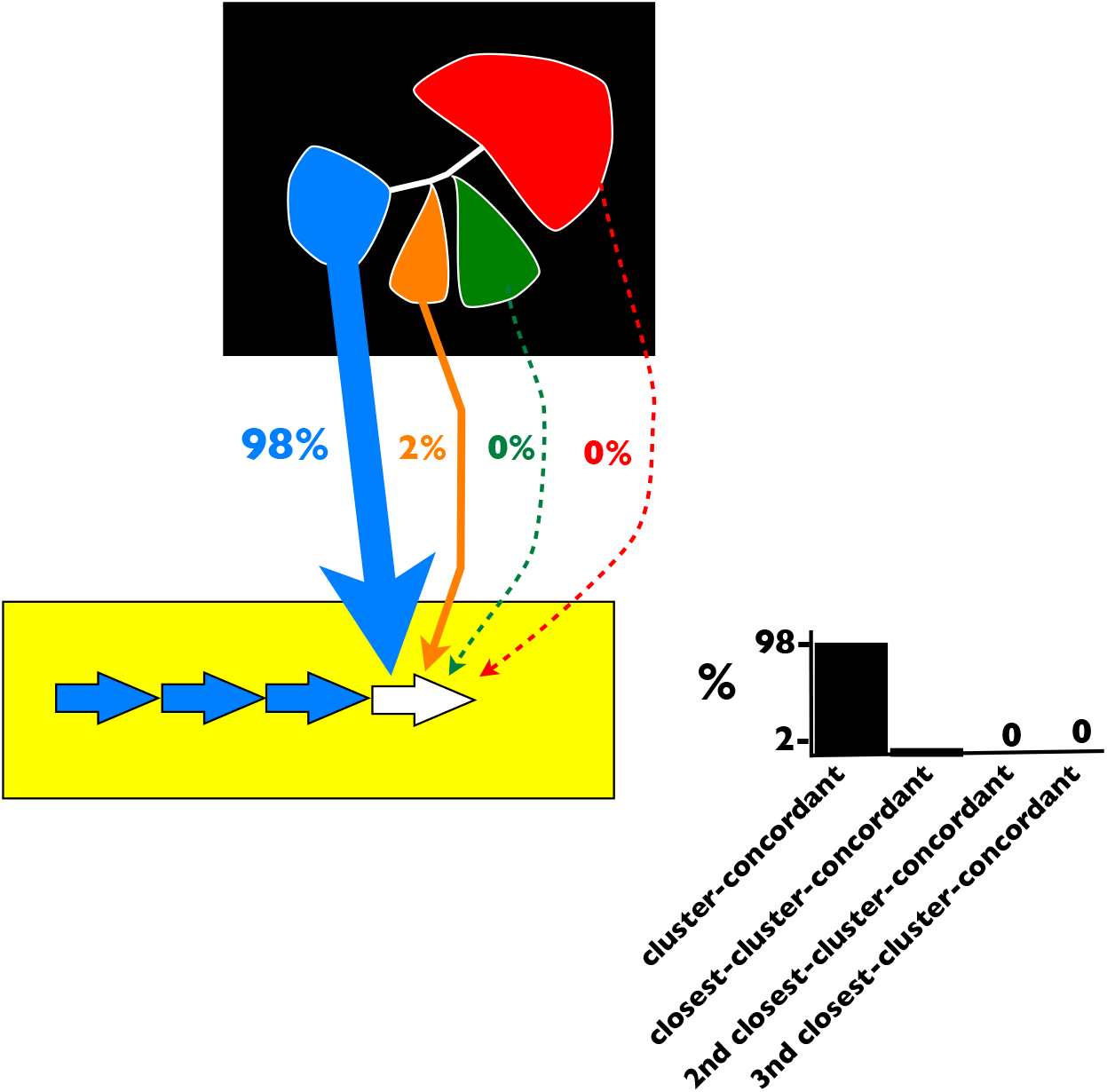
The pattern shown in Figure 6 of the main text suggests that if a new dimer is added to an existing HOR (white arrow), homology with extant dimers strongly influences this process.

**Supplemental Figure S18.**
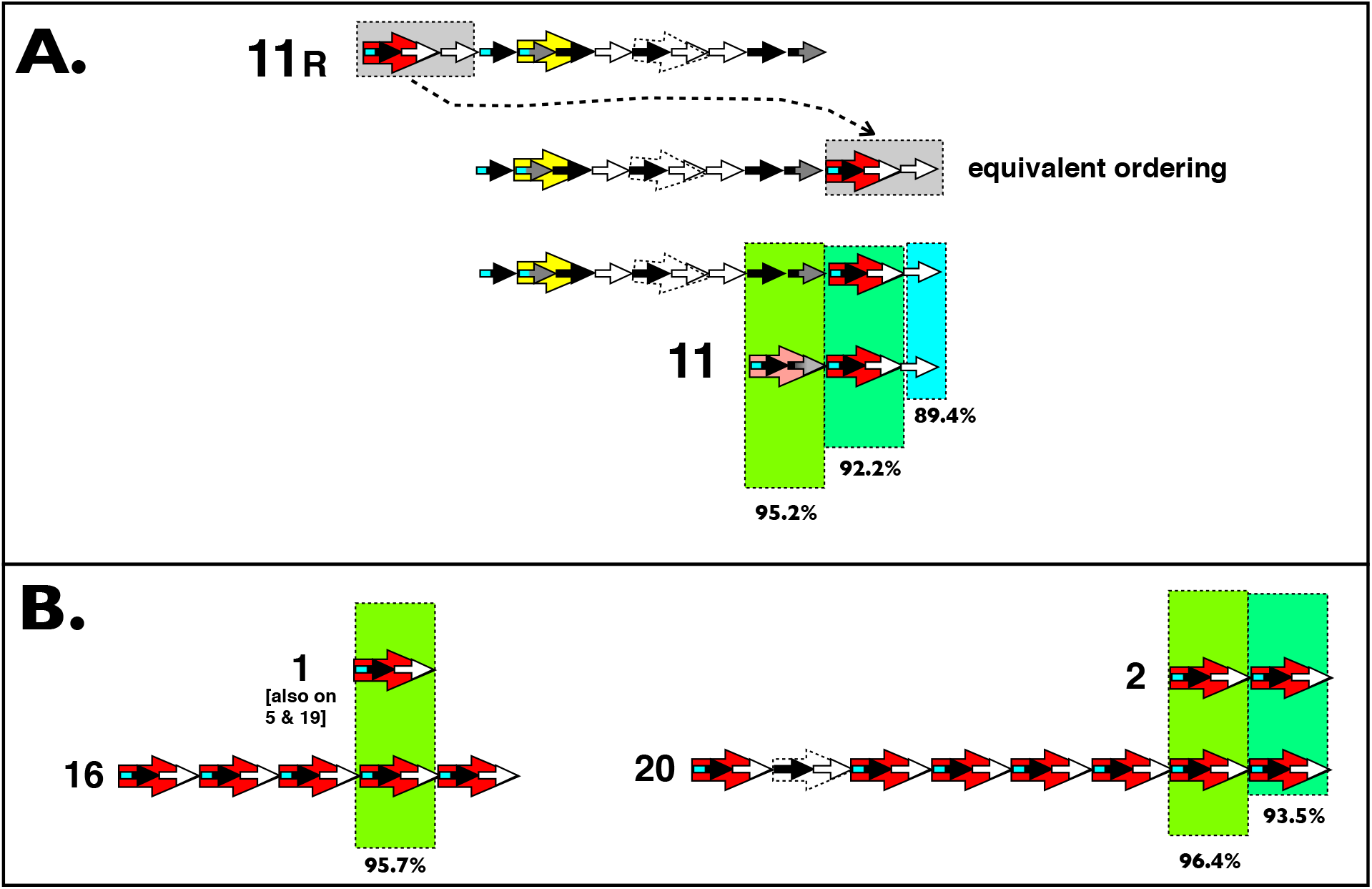
**A.** Top: schematic diagram of the consensus HOR immediately to the right of active HOR on chromosome 11 in the UCSC genome browser [GRCh38] (monomers are in the order shown in later parts of the paper [Figure 9]). To compare this inactive, flanking HOR to the active HOR, the first 3 monomers (grey box) have been moved to the end. The active HOR in chromosome 11 has high sequence similarity (percent values shown at bottom of figure) to a contiguous segment of the flanking, inactive HOR, and was feasibly generated via a contraction (deletion) within the inactive HOR. **B.** The smallest HORs (located on chromosome 1 [also on chromosomes 5 and 19] and chromosome 2) show close sequence similarity to dimers found on longer HORs on chromosomes 16 and 20.

**Supplemental Figure S19.**
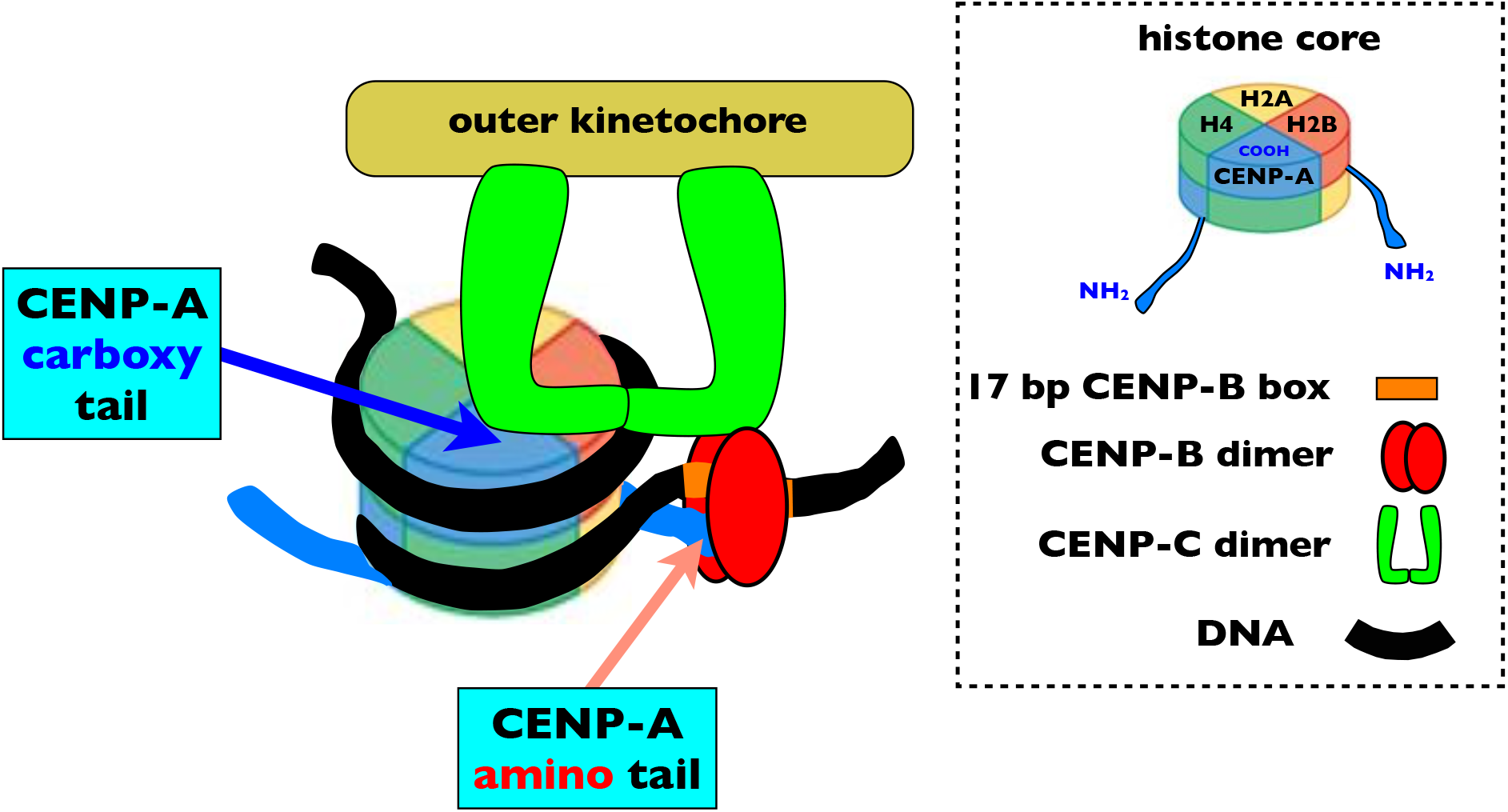
CENP-C binds to CENP-A-containing nucleosomes at two places: the carboxyl and amino ends of the molecule, and the attachment to the amino-end is dependent on the presence of CENP-B. Adapted from Fachinetti et al. (2015).

## References

Aldrup-MacDonald ME, Kuo ME, Sullivan LL, Chew K, Sullivan BA (2016) Genomic variation within alpha satellite DNA influences centromere location on human chromosomes with metastable epialleles. Genome research 26, 1301–1311.

Alkan, C., Cardone, M. F., Catacchio, C. R., Antonacci, F., O’Brien, S. J., Ryder, O. A., … & Ventura, M. (2011). Genome-wide characterization of centromeric satellites from multiple mammalian genomes. Genome research, 21(1), 137–145.

Arbeithuber, B., Betancourt, A. J., Ebner, T., & Tiemann-Boege, I. (2015). Crossovers are associated with mutation and biased gene conversion at recombination hotspots. Proceedings of the National Academy of Sciences, 112, 2109–2114.

Archidiacono N, Antonacci R, Marzella R, et al. (1995) Comparative mapping of human alphoid sequences in great apes using fluorescence in situ hybridization. Genomics 25, 477–484.

Aze, A., Sannino, V., Soffientini, P., Bachi, A., & Costanzo, V. (2016). Centromeric DNA replication reconstitution reveals DNA loops and ATR checkpoint suppression. Nature cell biology, 18(6), 684.

Bakhoum, S. F., Kabeche, L., Compton, D. A., Powell, S. N., & Bastians, H. (2017). Mitotic DNA damage response: At the crossroads of structural and numerical cancer chromosome instabilities. Trends in cancer, 3(3), 225–234.

Bhargava, R., Onyango, D. O., & Stark, J. M. (2016). Regulation of single-strand annealing and its role in genome maintenance. Trends in Genetics, 32(9), 566–575.

Black BE, Bassett EA (2008) The histone variant CENP-A and centromere specification. Current opinion in cell biology 20, 91–100.

Bodor DL, Mata JF, Sergeev M, et al. (2014) The quantitative architecture of centromeric chromatin. Elife 3, e02137.

Britten RJ (2002) Divergence between samples of chimpanzee and human DNA sequences is 5%, counting indels. Proceedings of the National Academy of Sciences 99, 13633–13635.

Bulazel, K., Metcalfe, C., Ferreri, G. C., Yu, J., Eldridge, M. D., & O’Neill, R. J. (2006). Cytogenetic and molecular evaluation of centromere-associated DNA sequences from a marsupial (Macropodidae: Macropus rufogriseus) X chromosome. Genetics, 172(2), 1129–1137.

Cam, H. P., Noma, K. I., Ebina, H., Levin, H. L., & Grewal, S. I. (2008). Host genome surveillance for retrotransposons by transposon-derived proteins. Nature, 451(7177), 431.

Cannan, W. J., & Pederson, D. S. (2016). Mechanisms and consequences of double-strand DNA break formation in chromatin. Journal of cellular physiology, 231(1), 3–14.

Casola, C., Hucks, D., & Feschotte, C. (2008). Convergent domestication of pogo-like transposases into centromere-binding proteins in fission yeast and mammals. Molecular biology and evolution, 25(1), 29–41.

Chan, K. L., & Hickson, I. D. (2011). New insights into the formation and resolution of ultra-fine anaphase bridges. In Seminars in cell & developmental biology (Vol. 22, No. 8, pp. 906–912). Academic Press.

Chaisson MJ, Wilson RK, Eichler EE (2015) Genetic variation and the de novo assembly of human genomes. Nature Reviews Genetics 16, 627–640.

Chakraborty, U., George, C. M., Lyndaker, A. M., & Alani, E. (2016). A delicate balance between repair and replication factors regulates recombination between divergent DNA sequences in Saccharomyces cerevisiae. Genetics, 202(2), 525–540.

Chmátal L, Gabriel SI, Mitsainas GP, et al. (2014) Centromere strength provides the cell biological basis for meiotic drive and karyotype evolution in mice. Current Biology 24, 2295–2300.

Choo, K. H., Vissel, B., Nagy, A., Earle, E., & Kalitsis, P. (1991). A survey of the genomic distribution of alpha satellite DNA on all the human chromosomes, and derivation of a new consensus sequence. Nucleic acids research, 19(6), 1179.

Costantino, L., Sotiriou, S. K., Rantala, J. K., Magin, S., Mladenov, E., Helleday, T., … & Halazonetis, T. D. (2014). Break-induced replication repair of damaged forks induces genomic duplications in human cells. Science, 343(6166), 88–91.

Crosetto, N., Mitra, A., Silva, M. J., Bienko, M., Dojer, N., Wang, Q., … & Pasero, P. (2013). Nucleotide-resolution DNA double-strand break mapping by next-generation sequencing. Nature methods, 10(4), 361.

Cui, P., Ding, F., Lin, Q., Zhang, L., Li, A., Zhang, Z., … & Yu, J. (2012). Distinct contributions of replication and transcription to mutation rate variation of human genomes. Genomics, proteomics & bioinformatics, 10(1), 4–10.

Durfy, S. J., & Willard, H. F. (1989). Patterns of intra-and interarray sequence variation in alpha satellite from the human X chromosome: evidence for short-range homogenization of tandemly repeated DNA sequences. Genomics, 5(4), 810–821.

Fachinetti D, Han JS, McMahon MA, et al. (2015) DNA sequence-specific binding of CENP-B enhances the fidelity of human centromere function. Developmental cell 33, 314–327.

Fishman L, Willis JH (2005) A novel meiotic drive locus almost completely distorts segregation in Mimulus (monkeyflower) hybrids. Genetics 169, 347–353.

Fishman-Lobell, J., Rudin, N., & Haber, J. E. (1992). Two alternative pathways of double-strand break repair that are kinetically separable and independently modulated. Molecular and cellular biology, 12(3), 1292–1303.

Fujita R, Otake K, Arimura Y, et al. (2015) Stable complex formation of CENP-B with the CENP-A nucleosome. Nucleic acids research 43, 4909–4922.

Fukagawa T, Earnshaw WC (2014) The centromere: chromatin foundation for the kinetochore machinery. Developmental cell 30, 496–508.

Gaff C, Du Sart D, KalltsIs P, et al. (1994) A novel nuclear protein binds centromeric alpha satellite DNA. Human molecular genetics 3, 711–716.

Greenfeder, S. A., & Newlon, C. S. (1992). Replication forks pause at yeast centromeres. Molecular and cellular biology, 12(9), 4056–4066.

Haaf, T., & Willard, H. F. (1992). Organization, polymorphism, and molecular cytogenetics of chromosome-specific α-satellite DNA from the centromere of chromosome 2. Genomics, 13(1), 122–128.

Haaf, T., Mater, A. G., Wienberg, J., & Ward, D. C. (1995). Presence and abundance of CENP-B box sequences in great ape subsets of primate-specific α-satellite DNA. Journal of molecular evolution, 41(4), 487–491.

Hasson D, Panchenko T, Salimian KJ, et al. (2013) The octamer is the major form of CENP-A nucleosomes at human centromeres. Nature structural & molecular biology 20, 687–695.

Hastings P (2010) Mechanisms of ectopic gene conversion. Genes 1, 427–439.

Hellborg, L., & Ellegren, H. (2004). Low levels of nucleotide diversity in mammalian Y chromosomes. Molecular Biology and Evolution, 21(1), 158–163.

Henikoff JG, Thakur J, Kasinathan S, Henikoff S (2015) A unique chromatin complex occupies young α-satellite arrays of human centromeres. Science advances 1, e1400234.

Henikoff S, Malik HS (2002) Centromeres: selfish drivers. Nature 417, 227–227.

Iwata-Otsubo A, Dawicki-McKenna JM, Akera T, et al. (2017) Expanded satellite repeats amplify a discrete CENP-A nucleosome assembly site on chromosomes that drive in female meiosis. Current Biology 27, 2365–2373. e2368.

Jeganathan, S., Petrovic, A., Singh, P., John, J., Krenn, V., Weissmann, F., … & Musacchio, A. (2016). Molecular basis of outer kinetochore assembly on CENP-T. Elife, 5, e21007.

Kipling, D., & Warburton, P. E. (1997). Centromeres, CENP-B and Tigger too. Trends in Genetics, 13(4), 141–145.

Kipling, D., Wilson, H. E., Mitchell, A. R., Taylor, B. A., & Cooke, H. J. (1994). Mouse centromere mapping using oligonucleotide probes that detect variants of them in or satellite. Chromosoma, 103(1), 46–55.

Kobayashi, T. (2014). Ribosomal RNA gene repeats, their stability and cellular senescence. Proceedings of the Japan Academy, Series B, 90(4), 119–129.

Komissarov, A. S., Gavrilova, E. V., Demin, S. J., Ishov, A. M., & Podgornaya, O. I. (2011). Tandemly repeated DNA families in the mouse genome. BMC genomics, 12(1), 531.

Kugou, K., Hirai, H., Masumoto, H., & Koga, A. (2016). Formation of functional CENP-B boxes at diverse locations in repeat units of centromeric DNA in New World monkeys. Scientific reports, 6, 27833.

Langley, S. A., Miga, K., Karpen, G. H., & Langley, C. H. (2018). Haplotypes spanning centromeric regions reveal persistence of large blocks of archaic DNA. BioRxiv.

Liu, Y., Nielsen, C. F., Yao, Q., & Hickson, I. D. (2014). The origins and processing of ultra fine anaphase DNA bridges. Current opinion in genetics & development, 26, 1–5.

Lo, A. W., Liao, G. C. C., Rocchi, M., & Choo, K. A. (1999). Extreme reduction of chromosome-specific α-satellite array is unusually common in human chromosome 21. Genome Research, 9(10), 895–908.

Malik HS, Henikoff S (2002) Conflict begets complexity: the evolution of centromeres. Current opinion in genetics & development 12, 711–718.

Maloney KA, Sullivan LL, Matheny JE, et al. (2012) Functional epialleles at an endogenous human centromere. Proceedings of the National Academy of Sciences 109, 13704–13709.

Marshall OJ, Chueh AC, Wong LH, Choo KA (2008) Neocentromeres: new insights into centromere structure, disease development, and karyotype evolution. The American Journal of Human Genetics 82, 261–282.

Masumoto H, Masukata H, Muro Y, Nozaki N, Okazaki T (1989) A human centromere antigen (CENP-B) interacts with a short specific sequence in alphoid DNA, a human centromeric satellite. The Journal of Cell Biology 109, 1963–1973.

Masumoto H, Yoda K, Ikeno M, et al. (1993) Properties of CENP-B and its target sequence in a satellite DNA. In: Chromosome segregation and aneuploidy, pp. 31–43. Springer.

McNulty, S. M., Sullivan, L. L., & Sullivan, B. A. (2017). Human centromeres produce chromosome-specific and array-specific alpha satellite transcripts that are complexed with CENP-A and CENP-C. Developmental cell, 42(3), 226–240.

McNulty, S. M., & Sullivan, B. A. (2018). Alpha satellite DNA biology: finding function in the recesses of the genome. Chromosome Research, 26(3), 115–138.

Miga KH, Newton Y, Jain M, et al. (2014) Centromere reference models for human chromosomes X and Y satellite arrays. Genome research 24, 697–707.

Mitra, S., Gomez-Raja, J., Larriba, G., Dubey, D. D., & Sanyal, K. (2014). Rad51–Rad52 mediated maintenance of centromeric chromatin in Candida albicans. PLoS genetics, 10(4), e1004344.

Muchová, V., Amiard, S., Mozgová, I., Dvořáčková, M., Gallego, M. E., White, C., & Fajkus, J. (2015). Homology-dependent repair is involved in 45 S rDNA loss in plant CAF-1 mutants. The Plant Journal, 81(2), 198–209.

Nechemia-Arbely Y, Fachinetti D, Miga KH, et al. (2017) Human centromeric CENP-A chromatin is a homotypic, octameric nucleosome at all cell cycle points. J Cell Biol 216, 607–621.

Ozenberger, B. A., & Roeder, G. S. (1991). A unique pathway of double-strand break repair operates in tandemly repeated genes. Molecular and Cellular Biology, 11 (3), 1222–1231.

Pâques, F., & Haber, J. E. (1999). Multiple pathways of recombination induced by double-strand breaks in Saccharomyces cerevisiae. Microbiol. Mol. Biol. Rev., 63(2), 349–404.

Pertile MD, Graham AN, Choo KA, Kalitsis P (2009) Rapid evolution of mouse Y centromere repeat DNA belies recent sequence stability. Genome research 19, 2202–2213.

Pironon N, Puechberty J, Roizès G (2010) Molecular and evolutionary characteristics of the fraction of human alpha satellite DNA associated with CENP-A at the centromeres of chromosomes 1, 5, 19, and 21. BMC genomics 11, 195.

Rhoads A, Au KF (2015) PacBio sequencing and its applications. Genomics, proteomics & bioinformatics 13, 278–289.

Rice WR (2019) A game of thrones at human centromeres II. a new molecular/evolutionary model. BioRxiv xxxxxx; doi: https://doi.org/xx.xxxx/xxxxxx.

Roizès G (2006) Human centromeric alphoid domains are periodically homogenized so that they vary substantially between homologues. Mechanism and implications for centromere functioning. Nucleic acids research 34, 1912–1924.

Romanova L, Deriagin G, Mashkova T, et al. (1996) Evidence for selection in evolution of alpha satellite DNA: the central role of CENP-B/pJα binding region. Elsevier.

Sakofsky CJ, Ayyar S, Malkova A (2012) Break-induced replication and genome stability. Biomolecules 2, 483–504.

Scott, K. C., & Bloom, K. S. (2014). Lessons learned from counting molecules: how to lure CENP-A into the kinetochore. Open biology, 4(12), 140191.

Shepelev, V. A., Alexandrov, A. A., Yurov, Y. B., & Alexandrov, I. A. (2009). The evolutionary origin of man can be traced in the layers of defunct ancestral alpha satellites flanking the active centromeres of human chromosomes. PLoS genetics, 5(9), e1000641.

Schildkraut, E., Miller, C. A., & Nickoloff, J. A. (2005). Gene conversion and deletion frequencies during double-strand break repair in human cells are controlled by the distance between direct repeats. Nucleic acids research, 33(5), 1574–1580.

Schueler, M. G., Dunn, J. M., Bird, C. P., Ross, M. T., Viggiano, L., Rocchi, M., Willard, HF, Eric D. Green, ED, and NISC Comparative Sequencing Program. (2005). Progressive proximal expansion of the primate X chromosome centromere. Proceedings of the National Academy of Sciences, 102(30), 10563–10568.

Schueler MG, Sullivan BA (2006) Structural and functional dynamics of human centromeric chromatin. Annu. Rev. Genomics Hum. Genet. 7, 301–313.

Scott, K. C. (2013). Transcription and ncRNAs: at the cent (rome) re of kinetochore assembly and maintenance. Chromosome research, 21(6-7), 643–651.

Seo J-S, Rhie A, Kim J, et al. (2016) De novo assembly and phasing of a Korean human genome. Nature.

Smith GP (1976) Evolution of repeated DNA sequences by unequal crossover. Science 191, 528–535.

Stephan W (1989) Tandem-repetitive noncoding DNA: forms and forces. Molecular biology and evolution 6, 198–212.

Stephan, W., & Cho, S. (1994). Possible role of natural selection in the formation of tandem-repetitive noncoding DNA. Genetics, 136(1), 333–341.

Sullivan, B. A., & Karpen, G. H. (2004). Centromeric chromatin exhibits a histone modification pattern that is distinct from both euchromatin and heterochromatin. Nature structural & molecular biology, 11(11), 1076.

Sullivan, L. L., Boivin, C. D., Mravinac, B., Song, I. Y., & Sullivan, B. A. (2011). Genomic size of CENP-A domain is proportional to total alpha satellite array size at human centromeres and expands in cancer cells. Chromosome research, 19(4), 457.

Syeda, A. H., Hawkins, M., & McGlynn, P. (2014). Recombination and replication. Cold Spring Harbor perspectives in biology, 6(11), a016550.

Talbert PB, Henikoff S (2010) Centromeres convert but don’t cross. PLoS biology 8, e1000326.

Tanaka K, Chang HL, Kagami A, Watanabe Y (2009) CENP-C functions as a scaffold for effectors with essential kinetochore functions in mitosis and meiosis. Developmental cell 17, 334–343.

Tanaka Y, Tachiwana H, Yoda K, et al. (2005) Human centromere protein B induces translational positioning of nucleosomes on α-satellite sequences. Journal of Biological Chemistry 280, 41609–41618.

Tawaramoto, M. S., Park, S. Y., Tanaka, Y., Nureki, O., Kurumizaka, H., & Yokoyama, S. (2003). Crystal structure of the human centromere protein B (CENP-B) dimerization domain at 1.65-Å resolution. Journal of Biological Chemistry, 278(51), 51454–51461.

Thompson, J. D., Sylvester, J. E., Gonzalez, I. L., Costanzi, C. C., & Gillespie, D. (1989). Definition of a second dimeric subfamily of human α satellite DNA. Nucleic acids research, 17(7), 2769–2782.

Tsouroula, K., Furst, A., Rogier, M., Heyer, V., Maglott-Roth, A., Ferrand, A., … & Soutoglou, E. (2016). Temporal and spatial uncoupling of DNA double strand break repair pathways within mammalian heterochromatin. Molecular cell, 63(2), 293–305.

Wang, L. H. C., Mayer, B., Stemmann, O., & Nigg, E. A. (2010). Centromere DNA decatenation depends on cohesin removal and is required for mammalian cell division. J Cell Sci, 123(5), 806–813.

Warmerdam, D. O., van den Berg, J., & Medema, R. H. (2016). Breaks in the 45S rDNA lead to recombination-mediated loss of repeats. Cell reports, 14(11), 2519–2527.

Waterson, G. A. (1975). On the number of segregating sites in genetical models without recombination. Theor. Popul. Biol, 7, 256–276.

Waye, J. S., & Willard, H. F. (1986A). Molecular analysis of a deletion polymorphism in alpha satellite of human chromosome 17: evidence for homologous unequal crossing-over and subsequent fixation. Nucleic acids research, 14(17), 6915–6927.

Waye, J. S., & Willard, H. F. (1986B). Structure, organization, and sequence of alpha satellite DNA from human chromosome 17: evidence for evolution by unequal crossing-over and an ancestral pentamer repeat shared with the human X chromosome. Molecular and cellular biology, 6(9), 3156–3165.

Waye, J. S., Creeper, L. A., & Willard, H. F. (1987A). Organization and evolution of alpha satellite DNA from human chromosome 11. Chromosoma, 95(3), 182–188.

Waye, J. S., England, S. B., & Willard, H. F. (1987B). Genomic organization of alpha satellite DNA on human chromosome 7: evidence for two distinct alphoid domains on a single chromosome. Molecular and cellular biology, 7(1), 349–356.

Willard HF, Waye JS (1987) Hierarchical order in chromosome-specific human alpha satellite DNA. Trends in Genetics 3, 192–198.

Willard, H. F. (1991). Evolution of alpha satellite. Current opinion in genetics & development, 1(4), 509–514.

Wilson Sayres, M. A., & Makova, K. D. (2011). Genome analyses substantiate male mutation bias in many species. Bioessays, 33(12), 938–945.

Wilson Sayres, M. A., Lohmueller, K. E., & Nielsen, R. (2014). Natural selection reduced diversity on human Y chromosomes. PLoS genetics, 10(1), e1004064.

Wong, A. K. C., & Rattner, J. B. (1988). Sequence organization and cytological localization of the minor satellite of mouse. Nucleic acids research, 16(24), 11645–11661.

Yang, J. W., Pendon, C., Yang, J., Haywood, N., Chand, A., & Brown, W. R. A. (2000). Human mini-chromosomes with minimal centromeres. Human Molecular Genetics, 9(12), 1891–1902.

Yoda, K., Kitagawa, K., Masumoto, H., Muro, Y., & Okazaki, T. (1992). A human centromere protein, CENP-B, has a DNA binding domain containing four potential alpha helices at the NH2 terminus, which is separable from dimerizing activity. The Journal of cell biology, 119(6), 1413–1427.

Yoda, K., Nakamura, T., Masumoto, H., Suzuki, N., Kitagawa, K., Nakano, M., … & Okazaki, T. (1996). Centromere protein B of African green monkey cells: gene structure, cellular expression, and centromeric localization. Molecular and cellular biology, 16(9), 5169–5177.

Zaratiegui, M., Vaughn, M. W., Irvine, D. V., Goto, D., Watt, S., Bähler, J., … & Martienssen, R. A. (2011). CENP-B preserves genome integrity at replication forks paused by retrotransposon LTR. Nature, 469(7328), 112.

Ziccardi W, Zhao C, Shepelev V, et al. (2016) Clusters of alpha satellite on human chromosome 21 are dispersed far onto the short arm and lack ancient layers. Chromosome research 24, 421–436.

